# Constitutive PDGFRβ activation drives connective tissue overgrowth through STAT5-IGF1 signaling

**DOI:** 10.64898/2026.08.27.747555

**Authors:** Hae Ryong Kwon, Alex Rackley, Lorin E. Olson

**Affiliations:** Cardiovascular Biology Research Program, Oklahoma Medical Research Foundation, Oklahoma City, OK 73104, USA

**Author notes:** Correspondence to: Lorin E. Olson, 825 NE 13^th^ Street, Oklahoma City, OK 73104.

**Keywords:** connective tissue, bone, cartilage, skin, overgrowth, fibrosis, Kosaki overgrowth syndrome, platelet-derived growth factor receptor beta, lineage tracing, signal transducer and activator of transcription 5, insulin-like growth factor 1, growth hormone

## Abstract

Autosomal dominant gain-of-function mutations in platelet-derived growth factor receptor beta (PDGFRβ) cause overgrowth of the skeleton and other connective tissue in Kosaki overgrowth syndrome. However, the target cell type and signaling pathways underlying PDGFRβ-driven overgrowth are unknown. Normal postnatal growth is controlled by pituitary-secreted growth hormone (GH), which activates the STAT5 transcriptional factor to upregulate insulin-like growth factor 1 (IGF1). To investigate the role of the GH-STAT5-IGF1 pathway in PDGFRβ-related overgrowth, we generated mice with a PDGFRβ gain-of-function mutation in skeletal and fibroblast lineages, which resulted in STAT5 activation and gigantism. Conditional deletion of *Stat5ab* in connective tissue lineages rescued skeletal overgrowth and keloid-like fibrosis in the skin. Conditional deletion of GH receptor (*Ghr*) did not rescue overgrowth, indicating the physiological activator of STAT5 is not required for overgrowth. However, deletion of *Igf1*, the STAT5 target gene, and its receptor, *Igf1r*, in connective tissue, rescued the overgrowth phenotype. These findings demonstrate a GHR-independent STAT5-IGF1 signaling pathway in mutant connective tissue cells, which mediates PDGFRβ-driven overgrowth in mice and potentially in humans with similar *PDGFRB* mutations.

## INTRODUCTION

Kosaki overgrowth syndrome (KOGS) is a rare genetic disorder caused by autosomal dominant germline mutations in platelet-derived growth factor receptor beta (PDGFRβ) that result in constitutive receptor activation. Clinical features include tall stature and coarse facial features, with progressive connective tissue changes affecting the skeleton (craniosynostosis, scoliosis, osteopenia), skin (thin, hyperelastic skin and hypertrophic scars), and vasculature (aneurysms, heart valve thickening)^1,2^. Penttinen Syndrome is also caused by autosomal dominant mutations in PDGFRβ and some PS patients exhibit tall stature like KOGS^3–5^. A third PDGFRβ-related disease is myofibromatosis (MF), caused by somatic gain-of-function mutations and characterized by connective tissue tumors^6^. Thus, constitutive activation of PDGFRβ is associated with diseases affecting connective tissue growth and homeostasis^7–10^. However, the underlying molecular pathogenesis is currently unclear.

Platelet-derived growth factor (PDGF) is an essential secreted signal that stimulates mesenchymal cells to proliferate and secrete ECM during development and tissue repair^11–13^. PDGFRβ is one of two mammalian receptors that bind extracellular PDGF ligands and activate intracellular tyrosine kinase activity. We previously generated mice to express PDGFRβ with a mutation (D849V, designated as K) that confers constitutive tyrosine kinase activation^14^. Heterozygous *Pdgfrb^K/+^* mice do not exhibit overgrowth but instead develop lethal autoinflammation mediated by signal transducer and activator of transcription 1 (STAT1)^14^. It was subsequently shown, however, that *Pdgfrb^K/+^* mice on a *Stat1^-/-^*genetic background are rescued from autoinflammation and instead develop connective tissue overgrowth dramatically affecting skeleton and skin^15^. This mouse phenotype resembled KOGS, although different gain-of-function PDGFRβ variants have been associated with human KOGS. In any case, the signaling mediator(s) of PDGFRβ-induced overgrowth remain unknown. We previously identified hyperactive STAT5 and elevated insulin-like growth factor 1 (IGF1) in skeletal stem cells from *Pdgfrb^K/+^*mice^16^. This led to the hypothesis that STAT5-IGF1 signaling may be involved in overgrowth.

Pituitary-secreted GH promotes postnatal growth via the STAT5 transcription factor, which regulates local and systemic IGF1 expression that ultimately drives growth of the skeleton, connective tissue, and most other cells in the body. Here, we used a genetic approach to test whether STAT5-IGF1 signaling mediates tissue overgrowth caused by overactive PDGFRβ, and the roles of growth hormone receptor (GHR) and IGF1 receptor (IGF1R). STAT5 consists of two distinct genes, *Stat5a* and *Stat5b,* encoding highly similar proteins that mediate signaling by a variety of cytokine receptors. Downstream of GHR, STAT5 controls the expression of IGF1. Secreted IGF1 binds to IGF1R, a receptor tyrosine kinase that promotes cell proliferation and tissue growth^17^. We used two different approaches for fibroblast and skeletal cell expression of *Pdgfrb^K^* and *Stat1-*deletion, which both resulted in overgrowth. We performed genetic epistasis experiments by deletion of *Stat5a/b, Ghr, Igf1, and Igf1r*. We found that *Pdgfrb^K^* requires *Stat5a/b, Igf1* and *Igf1r* to cause overgrowth. Deletion of *Ghr*, however, has no effect on PDGFRβ-driven overgrowth, suggesting that gain-of-function PDGFRβ signaling hijacks the growth hormone pathway by directly phosphorylating STAT5.

## RESULTS

### Overgrowth of PDGFRβ^K^Stat1 mice rescued by deletion of STAT5

Mice heterozygous for a global *Pdgfrb^K^* mutation and *Stat1^-/-^* genetic background (*Pdgfrb^+/K^Stat1^-/-^Sox2Cre* mice, denoted *KS1^Sox2Cre^*) develop overgrowth. Deletion of *Stat1* from *Pdgfrb^K^* mice is required to alleviate lethal autoinflammation and negative feedback on PDGFRβ, although *Stat1* deletion on its own does not cause overgrowth ^18–20^. Both sexes of *bKS1^Sox2Cre^* mice are overgrown, with dramatically increased body weight, size of the skeleton, and skin thickness^15,16^. To design experiments for expression *Pdgfrb^K^* and deletion *Stat1* from bone and skin in a tissue-specific manner, we analyzed publicly available scRNA-seq datasets^21–23^ and noted the expression pattern of genes in the GH/IGF1 pathway that could be involved in overgrowth. In bone, the highest co-expression of *Pdgfrb* and *Igf1* was in osteoblast lineages (bone marrow stromal cells and osteoblasts) that also express *Pdgfra* and *Prrx1*. Lower levels of *Stat5a, Stat5b, Igf1r* and *Ghr* were expressed in all types of skeletal cells, suggesting capacity to respond to IGF1 or GH (**Supplementary Figure 1A-B**). In skin, the highest co-expression of *Pdgfrb* and *Igf1* was in fibroblast lineages that also express *Pdgfra* and *Prrx1*. Again, *Stat5a, Stat5b, Igf1r* and *Ghr* were expressed in all skin cell types, suggesting capacity to respond to IGF1 or GH Fibroblasts mapped to two subpopulations designated by *Lef1* (fibroblast-1) or *Col1a1* (fibroblast-2). *Igf1* and *Ghr* were highest in fibroblast-2, *Igf1r* was highest in fibroblast-1 (**Supplementary Figure 1C**). Vascular smooth muscle cells (VSMC) express high *Pdgfrb,* but they were ruled out as the source of overgrowth because *bKS1* mutants generated with a VSMC-specific Cre are not overgrown^24^.

We devised two genetic approaches to induce expression of *Pdgfrb^K^*and deletion of *Stat1* in specific cellular lineages. To target bones and connective tissues of the developing limb, we used *Prrx1Cre*, which is active in the limb bud mesenchyme starting at E10.5^25^. We generated a cohort of *Pdgfrb^K^Stat1^flox^Prrx1Cre* (*bKS1^Prrx1Cre^*) mice with control littermates that lacked Cre or *Pdgfrb^K^* (**Supplementary Figure 2A**). The *bKS1^Prrx1Cre^* mice appeared normal at birth, but both sexes developed overgrowth with age (**Figure 1A**). To target dermal fibroblasts in a temporally restricted manner, we used *PdgfraCreER*^26^. A cohort *Pdgfrb^K^Stat1^flox^PdgfraCreER* (*bKS1^PdgfraCreER^*) mutant mice and controls (**Supplementary Figure 2D**) received a single tamoxifen treatment at 2 weeks old. The *bKS1^PdgfraCreER^* mice appeared normal at weaning, but both sexes developed overgrowth with age (**Figure 1B**). There is overlap between the two Cre drivers, because *Prrx1Cre* targets the ventral dermis in addition to the limb bud mesenchyme, and *PdgfraCreER* targets osteoblasts in addition to fibroblasts in postnatal skin. However, somewhat different phenotypes were expected because of different timing of activation, different target cell profiles, and more mosaicism with inducible *PdgfraCreER*. The above results show that mesenchymal lineages targeted by *Prrx1Cre* and *PdgfraCreER* are the source of overgrowth caused by *Pdgfrb^K^*.

**Figure 1.**
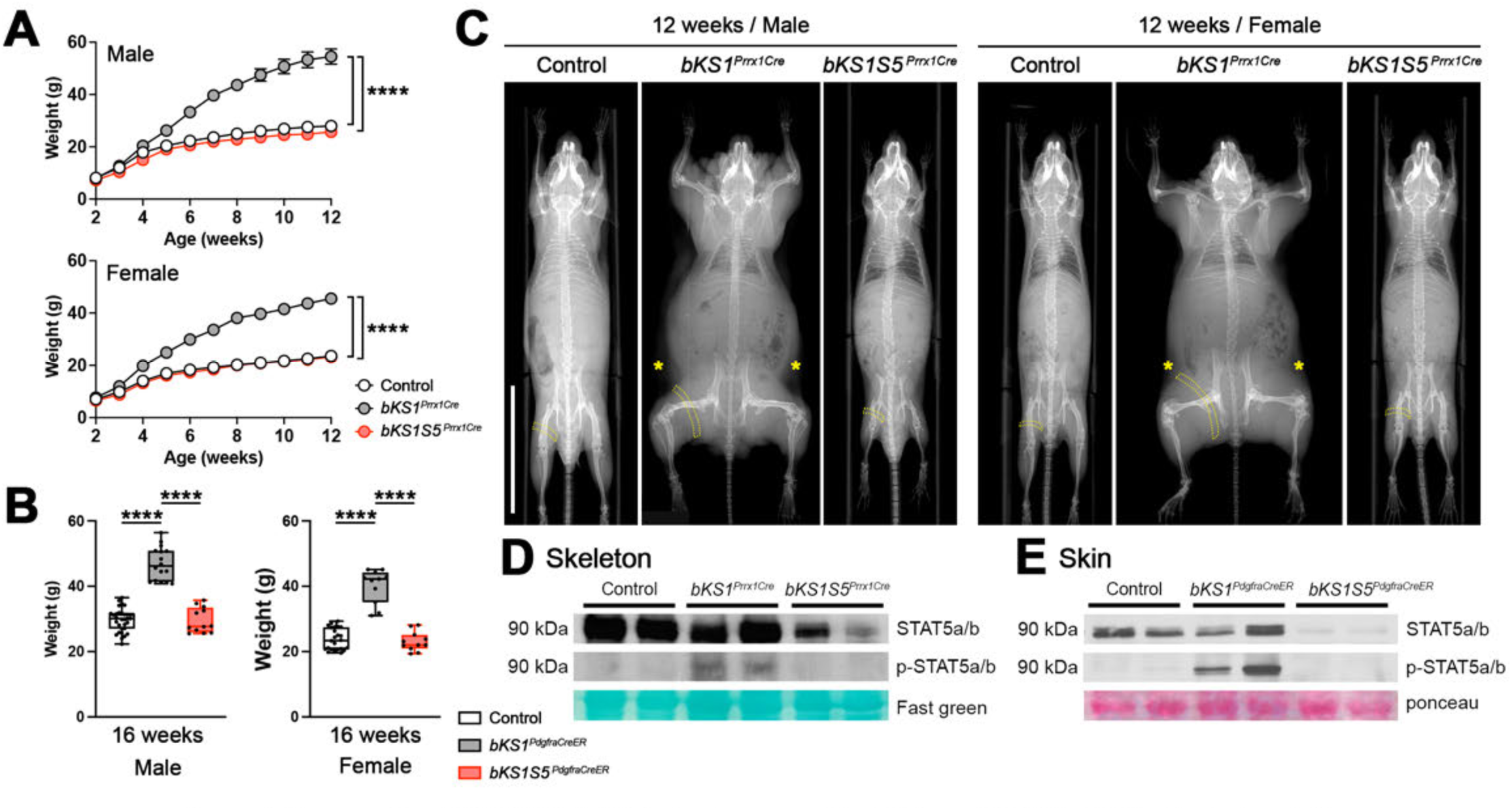
Overgrowth of PDGFRβ^K^Stat1^flox^ mice is rescued by deletion of STAT5. **(A)** Weekly body weight measurements of male and female control, *bKS1^Prrx1Cre^,* and *bKS1S5^Prrx1Cre^* mice from 2 to 12 weeks old. Male: n= 17 for control, n = 13 for *bKS1^Prnt1Cre^,* and n = 14 for *bKS1S5^Prrx1Cre^.* Female: n= 20 for control, n = 18 for *bKS1^Prrx1Cre^,* and n = 9 for *bKS1S5^PrrxlCre^.* **(B)** Final body weight measurements of male and female control, *bKS1^oRCreER^* and *bKS1S5^aRCreER^* mice at 16 weeks old. **(C)** X-ray images of male and female control, *bKS1^Prrx1Cre^* and *bKS1S5^Prrx1Cre^* mice at 12 weeks old. Dotted yellow bands indicate hindlimb muscle, hypertrophied in *bKS1^Prrx1Cre^.* Yellow asterisks indicate thick skin in *bKS1^Prrx1Cre^.* Scalebar, 3 cm. **(D-E)** Western blots of total and phosphorylated STAT5 in control, bKS1 and bKS1S5 in skeleton (femurs and tibias) (D) and skin (E). Fast green or ponceau stain shows similar protein loading. Statistical analyses are a mixed-effects model (A) and one-way ANOVA (B). ***, *p <* 0.0001.

To test the role of STAT5 in skeletal overgrowth, we introduced *Stat5a/b^flox^*alleles^27^ to generate a cohort of *bKS1S5^Prrx1Cre^*mice (**Supplementary Figure 2B**). A growth curve showed that *bKS1^Prrx1Cre^*weight began to diverge from the normal growth profile at 4-5 weeks of age, but *bKS1S5^Prrx1Cre^* mice of both sexes weighed the same as controls (**Figure 1A**). Whole body X-ray scan revealed an enlarged body and skeleton in male and female *bKS1^Prrx1Cre^* mice at 12 weeks. Thickening and elongation of the appendicular bones accompanied changes in joint structure that caused the limbs to rest in a bent position during scanning (**Figure 1C**). Overgrown mice were ambulatory in their cages, albeit with altered gait. Scans also highlighted muscle hypertrophy consistent with a previous study^28^ and skin thickening. In contrast, *bKS1S5^Prrx1Cre^* mice showed normalization of joint structure with a complete rescue of skin and muscle overgrowth (**Figure 1C**). We also generated *bKS1S5^PdgfraCreER^* mice with tamoxifen administered at 2 weeks of age (**Supplementary Figure 2E**). At 16 weeks, *bKS1S5^PdgfraCreER^* mice weighed the same as control mice while *bKS1^PdgfraCreER^* mice were significantly overgrown (**Figure 1B**). To determine the levels of STAT5, we isolated protein from hindlimb bones (*Prrx1Cre*) and skin (*PdgfraCreER*) for western blotting. As predicted from earlier work^15,16^, phosphorylated STAT5 was elevated in *bKS1* samples, but it was absent in *bKS1S5* samples (**Figure 1D-E**). We conclude that removal of activated STAT5 from *bKS1* fibroblasts and skeletal cells rescues overgrowth, demonstrating the required role of STAT5 for overgrowth.

### Bone and cartilage expansion depends on STAT5

Tibial length and cortical bone measurements showed that *bKS1^Prrx1Cre^*tibias were longer and thicker, but *bKS1S5^Prrx1Cre^* tibias were undergrown compared to controls (**Figure 2A-C**). Bone volume (BV) and total volume (TV) were increased in *bKS1^Prrx1Cre^* compared to controls but rescued in *bKS1S5^Prrx1Cre^* (**Figure 2C**). There were no changes in BV/TV ratio and mineral density. The growth plate is a major driver of long bone growth during puberty^29,30^. We examined growth plate structure and histomorphometry in 3-week-old proximal tibia. Compared to controls, *bKS1^Prrx1Cre^* growth plates were expanded in height and area, and the enthesis was more than twice as wide. These features were normalized in *bKS1S5^Prrx1Cre^* (**Figure 2D**). Within the growth plate, the resting and hypertrophic zones were significantly larger in *bKS1^Prrx1Cre^*mutants compared to controls, and *bKS1S5^Prrx1Cre^* reduced the proliferating and hypertrophic zones below the level of controls (**Figure 2E**). Therefore, bone and cartilage were expanded in the *bKS1^Prrx1Cre^* appendicular skeleton in a STAT5-dependent manner.

**Figure 2.**
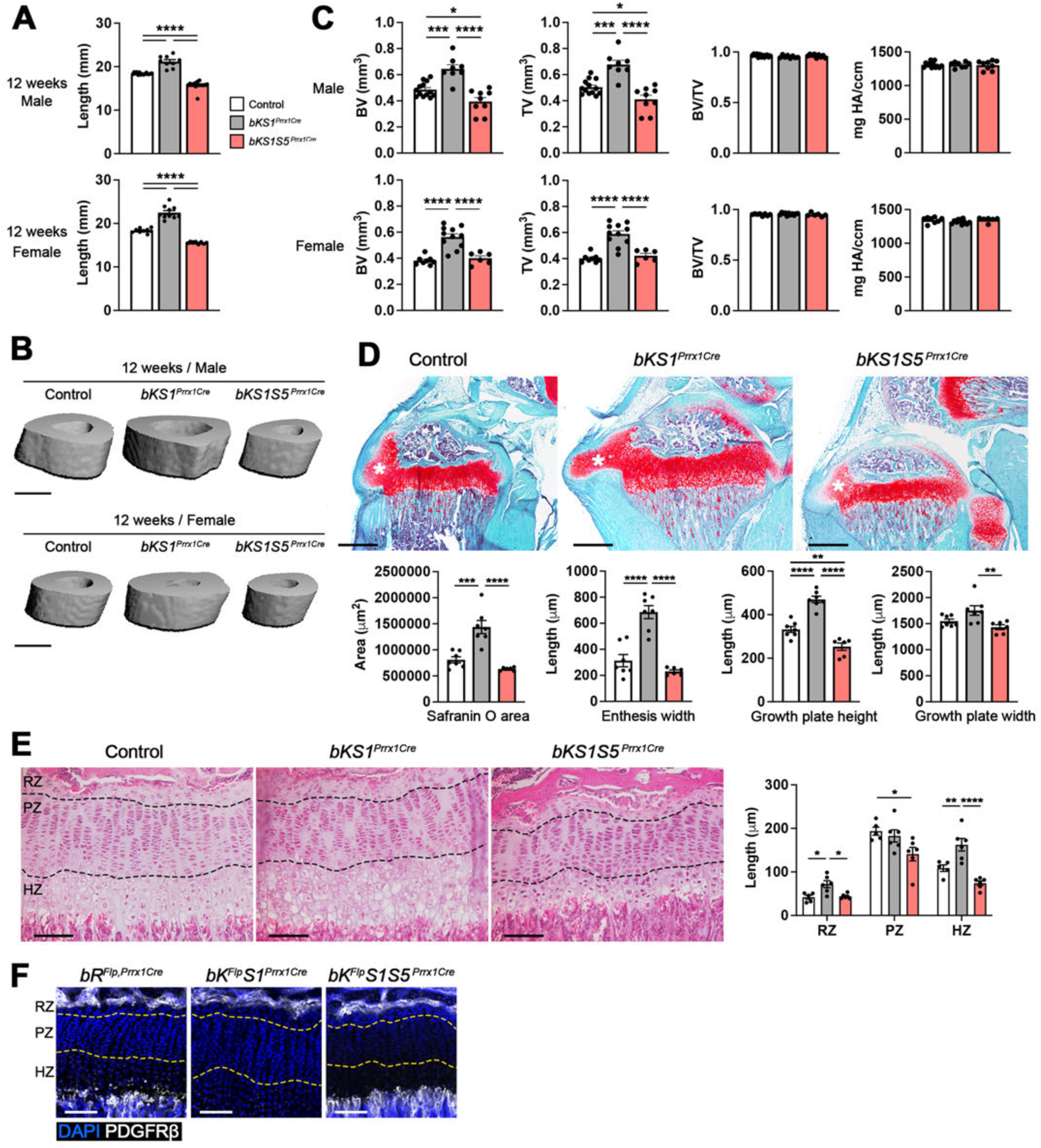
STAT5 deletion rescues tibia length and growth plate expansion. **(A)** Measurements of male and female tibia length from control, *bKS1^Prrx1Cre^,* and *bKS1 S5^Prrx1Cre^* mice at 12 weeks old. **(B-C)** Micro computed tomography scans and quantifications of male and female tibia cortical bone from control, *bKS1^Prrx1Cre^,* and *bKS1S5^Prrx1Cre^.* BV = bone volume, TV = total volume, BV/TV = bone volume ratio, mg HA/cm^3^ = mineral density. Scalebar, 0.5 cm. **(D)** Safranin O stain shows cartilage area in proximal tibias at 3 weeks old. Safranin O area, enthesis width, growth plate height, and growth plate width are quantified below. White asterisks indicate enthesis. Scalebar, 500 pm. **(E)** Hematoxylin and eosin stain of growth plates at 3 weeks old. Resting zone (RZ), proliferating zone (PZ), and hypertrophic zone (HZ) are quantified on the right. Scalebar, 100 pm. **(F)** Immunostaining of PDGFRp in tibia growth plate at 3 weeks old, with expression in the upper resting zone and in trabecular bone below. Scalebar, 100 pm. Statistical analyses are one-way ANOVA (A, C, D, and E). *, *p <* 0.05, **, *p <* 0.01, ***, *p <* 0.0005, ****, *p <* 0.0001.

Immunostaining showed that PDGFRβ was expressed by progenitor cells in the uppermost part of the resting zone but not in the proliferating and hypertrophic zones (**Figure 2F**). PDGFRβ is also highly expressed in periosteum and endosteum (**Supplementary Figure 3A**). To identify PDGFRβ^+^ skeletal progenitors and their progeny within the *Prrx1Cre* lineage, we performed intersectional lineage tracing with Flp and Cre. We used *Pdgfrb^Flp^*mice with a cassette containing a floxed cDNA encoding *Pdgfrb* followed by a *Flp°* sequence, which was inserted to replace the endogenous *Pdgfrb* gene. Cre recombination leads to expression of Flp° controlled by the *Pdgfrb* gene^31^. Thus, intersectional tracing can be achieved by combining *Prrx1Cre* with *Pdgfrb^Flp^* and a Flp-inducible R26-FSF-tdTomato reporter, resulting in *bR^Flp,Prrx1Cre^* mice with wild-type phenotype. In these mice, any Prrx1-derived cells that express *Pdgfrb^Flp^* will acquire indelible Tomato labeling **(Figure 3A**). In 3-week-old *bR^Flp,Prrx1Cre^*mice, all bone lineages and synovial joint structures were strongly Tomato^+^. Articular and enthesis cartilage showed partial labeling (**Figure 3C, F**). Chondrocytes were barely labeled at 3-week-old, in agreement with immunostaining. Tomato^+^ cells in the upper resting zone were sparse, and one or two chondrocyte columns were labeled per section (**Figure 3G**). By 6-months, however, many chondrocyte columns were labeled in *bR^Flp,Prrx1Cre^* (**Supplementary Figure 3B**). Therefore, PDGFRβ-lineage progenitors are not a major source of growth plate chondrocytes in young wild type mice.

**Figure 3.**
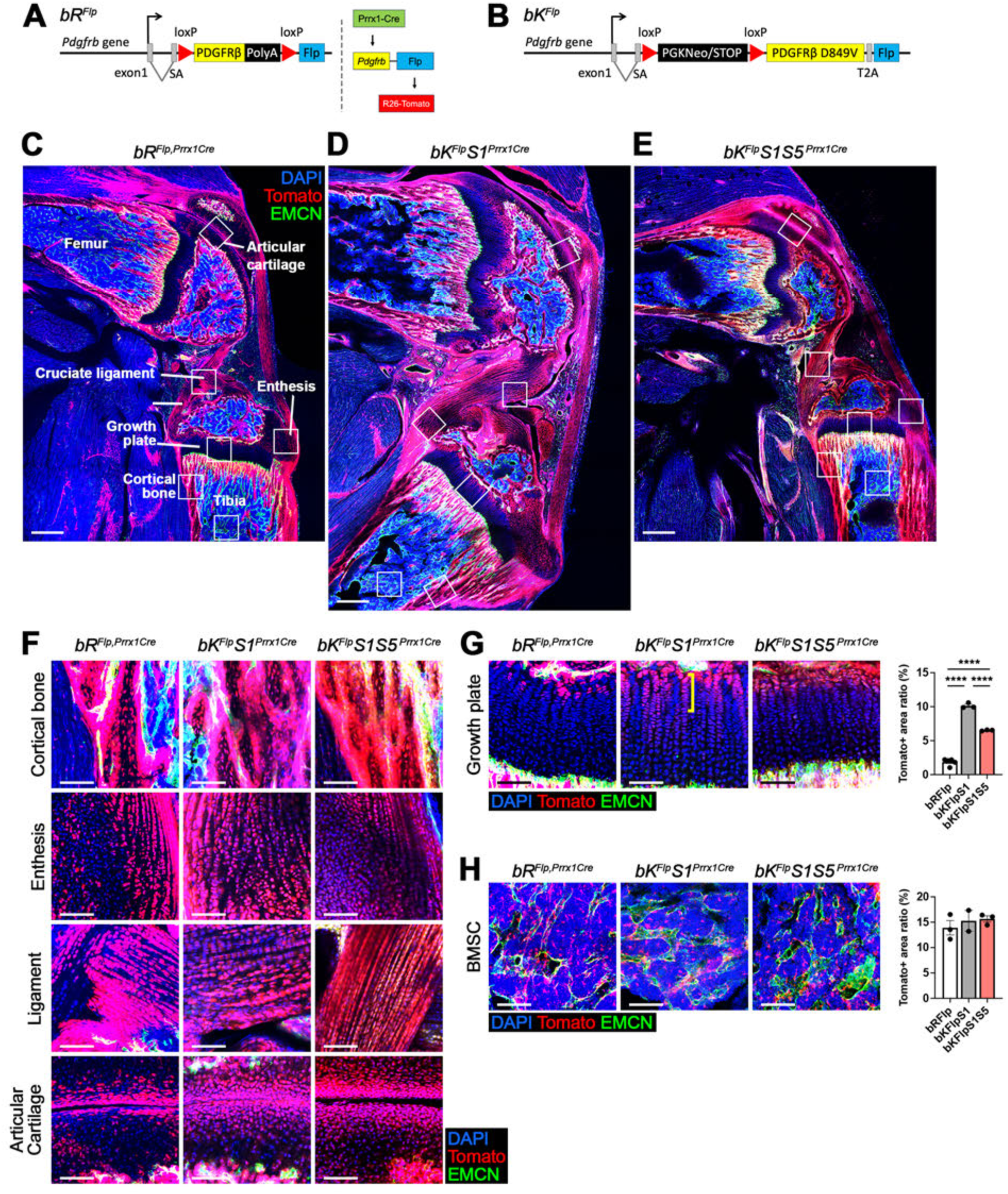
Intersectional lineage tracing of PDGFRβ in *Prrx1-Cre* skeleton. **(A)** Scheme of bR^Flp^ knock-in construct, containing splice acceptor (SA), loxP, wild type PDGFRβ cDNA and STOP, loxP, and Flp° recombinase. This construct allows conditional Flp° expression from one *Pdgfrb* allele. Diagram to the right shows the logic of intersectional lineage tracing: Cre recombinase triggers Flp expression under control of *Pdgfrb.* Then Flp° recombinase triggers the Tomato lineage reporter. **(B)** Scheme of bK^Flp^ knock-in construct, containing splice acceptor, lox-STOP-lox, PDGFRβ^K^ cDNA, T2A sequence, and Flp° recombinase. This construct allows conditional expression of PDGFRβ^K^ and Flp° from one *Pdgfrb* allele. **(C-E)** Confocal microscopy of distal femur, joint, and proximal tibia of *bR^Flp,Prrx1Cre^* (C), *bK^Flp^S1^Prrx1Cre^* (D), and *bK^Flp^S1S5^Prrx1Cre^* (E) at 3 weeks old. Tomato labeling indicates cells expressing bR^Flp^ or bK^Flp^. Endomucin (EMCN) stains vascular endothelial cells. Scalebar, 500 pm. (F-H) Zoomed images of cortical bone, enthesis, cruciate ligament, articular cartilage (F), growth plate (G) and bone marrow stromal cells (H) from C, D, and E images (insets). Tomato area ratio was measured in growth plate (G) and BMSC (H). A yellow bracket in growth plate (G) indicates an example of Tomato* cartilage column. Scalebar, 100 pm. Statistical analyses are one-way ANOVA. **“, *p <* 0.0001.

To perform intersectional tracing in the context of gain-of-function PDGFRβ signaling, we generated a new Cre-inducible *Pdgfrb^KFlp^* allele^32^. In these mice, Cre recombination leads to deletion of a Stop cassette to allow expression of the *Pdgfrb^K^* cDNA and Flp° (**Figure 3B**). We combined this new allele with *Prrx1Cre* and *Stat1^flox^* to create *bK^Flp^S1^Prrx1Cre^* mice with the same overgrowth phenotype as *bKS1^Prrx1Cre^*. In 3-week-old *bK^Flp^S1^Prrx1Cre^*mice, Tomato^+^ cells were abundant in the enlarged cortical bone, enthesis, ligament and other synovial joint structures, much like control mice (**Figure 3D, F**). There were significantly more columns of Tomato^+^ chondrocytes in *bK^Flp^S1^Prrx1Cre^* compared to controls (**Figure 3G**), but most columns remained unlabeled. This suggests that growth plate expansion may be indirectly influenced by effects of PDGFRβ^K^ in surrounding tissues. In bone marrow stromal cells (BMSCs), Tomato-labeling was similar between controls and *bK^Flp^S1^Prrx1Cre^* (**Figure 3H**).

We introduced *Stat5a/b^flox^* alleles to generate *bK^Flp^S1S5^Prrx1Cre^*mice for intersectional tracing. The resulting Tomato^+^ cells were distributed in bone and synovial joint tissues like controls and *bK^Flp^S1^Prrx1Cre^* (**Figure 3E-F**). Growth plate chondrocyte labeling was partially reduced in *bK^Flp^S1S5^Prrx1Cre^*compared to *bK^Flp^S1^Prrx1Cre^* but was still significantly higher than controls (**Figure 3G**). Taken together, these data suggest that PDGFRβ^K^-STAT5 signaling is active in bone, synovial, and some cartilage cell lineages, but indirect mechanisms emanating from PDGFRβ^+^ cells lead to growth plate expansion.

### Keloid-like dermal fibrosis depends on STAT5

Adult mouse dermis includes three layers: an upper layer of papillary dermis required for hair follicle formation, a middle layer of reticular dermis enriched for fibrillar collagen, and a bottom layer of dermal white adipose tissue (dWAT) defined by adipocytes^33,34^. We analyzed *bKS1^PdgfraCreER^*skin at 16 weeks to characterize phenotypic changes compared to controls. Trichrome stain revealed that male *bKS1^PdgfraCreER^* skin was abnormally thick with an expanded reticular dermis that obliterated the dWAT as a histologically distinct tissue layer (**Figure 4A**). Male *bKS1^PdgfraCreER^* skin was dramatically thicker than control skin due to expansion of reticular dermis, with obliteration of the dWAT in every case (**Figure 4B**). Picrosirus red stain revealed dense collagen bundles of abnormally large size reminiscent of keloid collagen ^35,36^ (**Figure 4C**). Female *bKS1^PdgfraCreER^* skin exhibited milder fibrosis with preservation of a distinct dWAT layer albeit with increased collagen deposition between adipocytes (**Supplementary Figure 4A**). Female *bKS1^PdgfraCreER^*overall skin thickness and dWAT thickness were not changed compared to control skin, but *bKS1^PdgfraCreER^* reticular dermis was slightly thicker than controls (**Supplementary Figure 4B**). Remarkably, knockout of *Stat5a/b* completely rescued the skin phenotype in both sexes, resulting in *bKS1S5^PdgfraCreER^* skin that was histologically and quantitatively indistinguishable from controls (**Figure 4A-C, Supplementary Figure 4A-B**). These results characterize *bKS1^PdgfraCreER^*skin overgrowth as fibrotic, STAT5-dependent, and sexually dimorphic.

**Figure 4.**
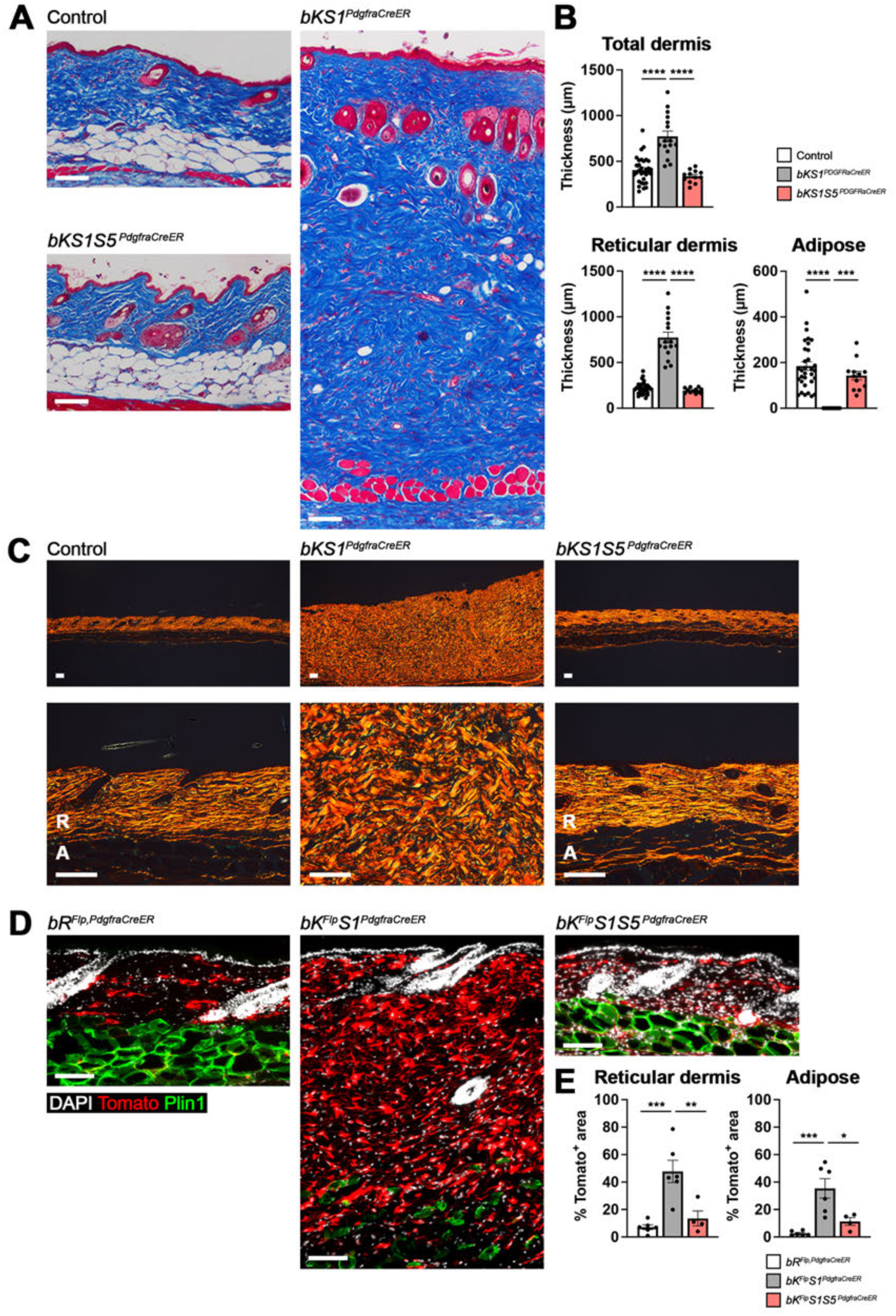
Keloid-like dermal fibrosis depends on STAT5. **(A-B)** Trichrome stain of male skin from control, *bKS1^aRCraER^,* and *bKS1 S5^aRCraER^* mice at 16 weeks old. Quantifications of thickness of total dermis, reticular dermis, and dermal white adipose tissue are shown. Scalebar, 100 pm. **(C)** Picrosirius red stain of male skin at 16 weeks old, R = reticular dermis, A = adipose. Scalebar, 100 pm. **(D-E)** Fluorescent images of male skins stained for adipocyte marker Perilipinl (Plinl, green). Tomato labeling indicates intersectional lineage tracing of *PdgfraCreER* and *bR^Flp^or bK^Flp^.* Scalebar, 100 pm. Quantification ofTomato* area in reticular dermis and dermal white adipose tissue are shown. Statistical analyses are one-way ANOVA (A and C). *, *p* < 0.05, **, p < 0.01, ***, p< 0.0005, ****,p< 0.0001.

We performed intersectional lineage tracing to identify PDGFRβ^+^ dermal cells and their progeny within the *PdgfraCreER* lineage. In *bR^Flp, PdgfraCreER^* skin with normal PDGFRβ signaling, Tomato^+^ cells were abundant in the reticular dermis and sparse in the papillary dermis and dWAT (**Figure 4D**). In male *bK^Flp^S1^PdgfraCreER^*skin, the expanded reticular dermis was composed of Tomato^+^ cells at high density, and Tomato^+^ cells filled the space between shrunken adipocytes (**Figure 4D**). Tomato^+^ cells were negative for the adipocyte marker Plin1 (**Figure 4D**) and expressed vimentin, consistent with fibroblast identity (**Supplementary Figure 4E**). In *bR^Flp, PdgfraCreER^* skin, Tomato^+^ fibroblasts occupied ∼7% of total area in the upper dermis and ∼2.5% in the lower dermis. In male *bK^Flp^S1^PdgfraCreER^* skin, Tomato^+^ fibroblasts increased to 47% and 35% of total area in the reticular dermis and lower dermis where dWAT should be (**Figure 4E**). In female *bK^Flp^S1^PdgfraCreER^* skin, the increase in Tomato^+^ fibroblast density was only significant for the dWAT (**Supplementary Figure 4C-D**). Deletion of *Stat5a/b* reversed the expansion of Tomato^+^ fibroblasts in both sexes (**Figure 4D-E, Supplementary Figure 4C-D**).

### Gene expression changes in overgrown skeleton and skin were rescued by STAT5 deletion

To gain insight into gene expression changes in skeleton, we performed bulk RNA sequencing with mRNA isolated from the tibia of control, *bKS1^Prrx1Cre^*, and *bKS1S5^Prrx1Cre^* mice at 6 weeks old. Compared to controls, there were 4885 differentially expressed genes (DEGs) in *bKS1^Prrx1Cre^*and 3054 in *bKS1S5^Prrx1Cre^*, with 1489 shared DEGs (**Figure 5A, Supplementary Figure 5A-B**). Among the shared DEGs, 49.6% showed reduced log_2_ fold change in *bKS1S5^Prrx1Cre^*compared to *bKS1^Prrx1Cre^,* indicating partial normalization of expression with *Stat5ab* deletion (**Supplementary Figure 5C**). Categorization of DEGs between control and *bKS1^Prrx1Cre^*identified upregulation of many bone-related gene ontologies including cartilage, collagen, ossification, ribosome, and insulin signaling (**Figure 5C, Supplementary Figure 5D**). Many individual DEGs were upregulated in overgrown skeleton and rescued by *Stat5ab* knockout, including major cartilage and bone regulatory genes *Sox9, Col2a1, Pthrp1* and *Spp1* (**Figure 5C, Supplementary Figure 6**). Importantly, the STAT5 target genes *Igf1* and *Socs2* were also upregulated in *bKS1^Prrx1Cre^* and normalized in *bKS1S5^Prrx1Cre^* tibias (**Figure 5B, Supplementary Figure 6C**).

**Figure 5.**
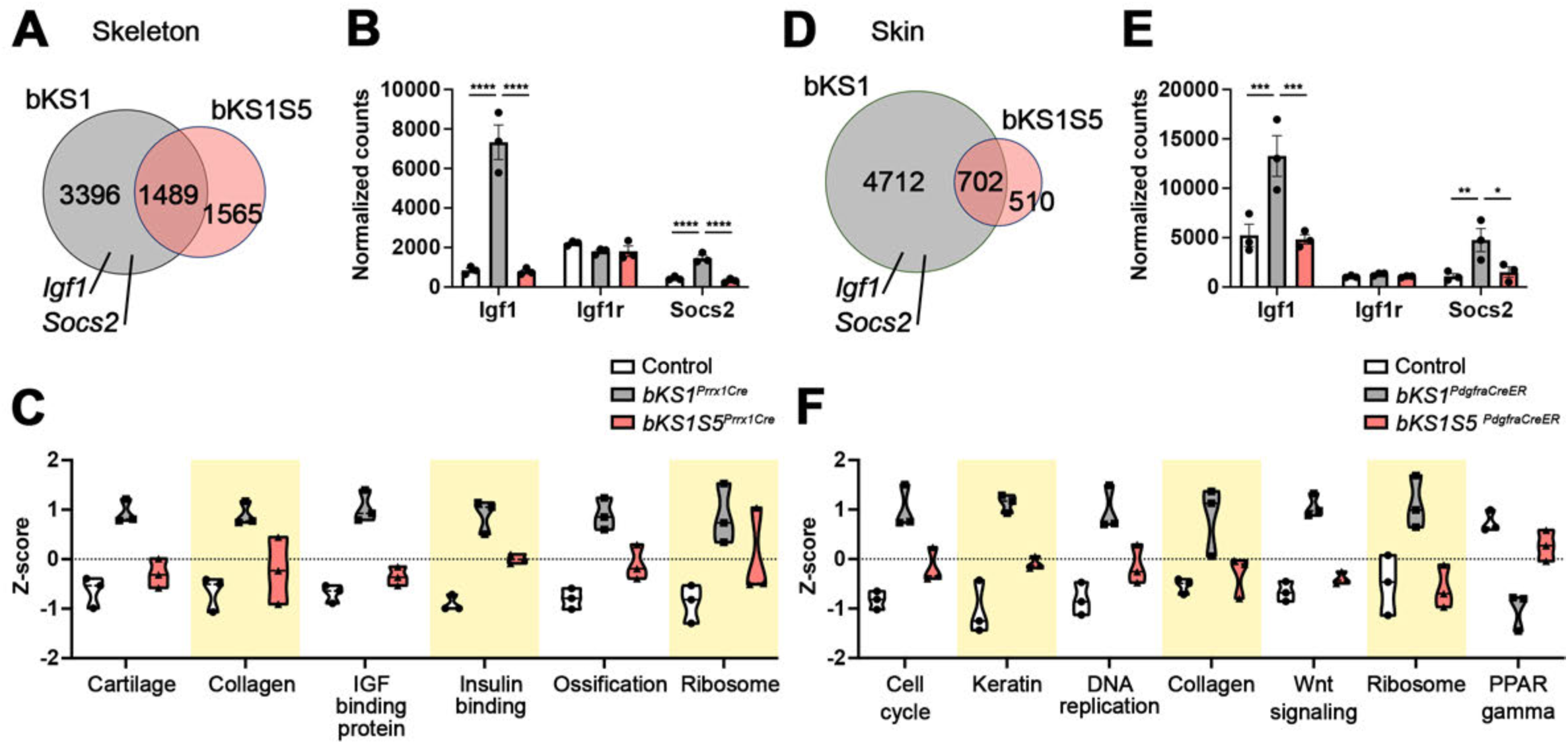
STAT5 deletion rescues gene expression changes in overgrown skeleton and skin. Tissue from control. bKS1. and bKS1 S5 mice was used for bulk RNA sequencing, n = 2 males and 1 female per genotype. **(A)** Venn Diagram showing *bKS1^Prrx1Cre^* specific, *bKS1S5^Prrx1Cre^* specific, and overlapping DEGs. **(B)** Normalized counts plots of *Igf1. Igflr,* and *Socs2* in control. bKS1, and bKS1S5 skeletons. **(C)** Quantification of gene-set expression across genotypes in skeletal tissues. Mean Z-scores were calculated across all genes (Supplementary Figure 6) within each gene category. (D) Venn Diagram showing *bKS1^Pd9fraCreER^* specific. *bKS1 S5^Pd9fraCreER^* specific, and overlapping DEGs. (E) Normalized counts plots of *Igf1, Igflr,* and *Socs2* in control, bKS1, and bKS1S5 skins. (F) Quantification of gene-set expression across genotypes in skin tissues. Mean Z- scores were calculated across all genes (Supplementary Figure 7 and 8) within each gene category. Statistical analyses (adjusted p-value) are Wald test in DESeq2 (B and D). *. *p* < 0.05, “. *p <* 0.01, ***, *p <* 0.0005, “**. *p* <0.0001.

For skin, we performed bulk RNA sequencing on mRNA isolated from control, *bKS1^PdgfraCreER^*, and *bKS1S5 ^PdgfraCreER^*skin at 16 weeks. Compared to controls, there were 5414 DEGs in *bKS1^PdgfraCreER^*and 1212 DEGs in *bKS1S5^PdgfraCreER^*, with 702 shared DEGs (**Figure 5D, Supplementary Figure 5E-F**). Among the shared DEGs, 88.2% showed reduced log_2_ fold change in *bKS1S5^PdgfraCreER^* compared to *bKS1^PdgfraCreER^*(**Supplementary Figure 5G**). Categorization of DEGs between control and *bKS1^PdgfraCreER^* identified upregulation of cell cycle, keratin, collagen, Wnt signaling, and PPARγ (**Figure 5F, Supplementary Figure 7**). Individual genes upregulated in *bKS1^PdgfraCreER^* and rescued in *bKS1^PdgfraCreER^* included the collagens *Col1a1, Col3a1*, Wnt signaling mediators *Axin1* and *Lef1*, and dozens of hair keratin-associated proteins (*Krtap*) (**Supplementary Figure 7**). PPARγ signaling as downregulated in *bKS1^PdgfraCreER^* and rescued in *bKS1S5^PdgfraCreER^* skin, consistent with change in dermal fat across the genotypes (**Supplementary Figure 5I**). Some categories were downregulated in *bKS1^PdgfraCreER^*and normalized in *bKS1S5^PdgfraCreER^*, including basement membrane, peroxisome, and mitochondrial genes (**Supplementary Figure 8**). Like the skeleton, *Igf1* and *Socs2* were upregulated in *bKS1^PdgfraCreER^* and normalized in *bKS1S5^PdgfraCreER^* skin (**Figure 5E**). Taken together, these results identify overgrowth- and fibrosis-related gene sets in the skin and skeleton, which are induced in *bKS1* and normalized in *bKS1S5*, thus demonstrating the functional significance of STAT5.

### Growth hormone receptor is not required for overgrowth

Growth hormone governs somatic growth control and sexually dimorphic growth rates, such that lack of circulating GH or loss of GH receptor diminishes STAT5 activation, reduces IGF1 levels, and produces dwarfism. The physiological effects of GH signaling defects become apparent at 3-4 weeks old in mice, which mirrors the time of overgrowth in *bKS1^Prrx1Cre^* mice (**Figure 1A**). Therefore, we were curious whether GHR might have a role in PDGFRβ-mediated overgrowth, perhaps supporting basal somatic growth that PDGFRβ^K^ mutation then amplifies. To test this idea, we introduced *Ghr^flox^* alleles^37^ to generate cohorts of *bKS1Ghr^Prrx1Cre^*and *bKS1Ghr^PdgfraCreER^* mice (**Supplementary Figure 2C, E**). We confirmed that *Ghr* produced a functionally null allele by generating dwarf mice upon homozygous germline mutation with *Sox2Cre* (**Supplementary Figure 9A**). However, *Ghr* deletion from *Prrx1*-lineage skeletal cells had no effect on PDGFRβ-mediated overgrowth, as *bKS1Ghr* males and females were the same as *bKS1* mutants in terms of weight and microCT scans (**Figure 6A-B, Supplementary Figure 9B-C**). Tibial lengths and cortical bone volumes were also not rescued (**Figure 6C-E, Supplementary Figure 9D-F**). Bone volume to total volume ratio was slightly decreased in *bKS1Ghr^Prrx1Cre^* males (**Figure 6E**). Interestingly, *Ghr* deletion from *Pdgfra*-lineage fibroblasts had a sexually dimorphic effect. Male weight was not rescued, but female weight was partially rescued (**Figure 6F, Supplementary Figure 9G**). Similarly, skin fibrosis and lipodystrophy were not rescued in *bKS1Ghr* males (**Figure 6G-H**). But inter-adipocyte fibrosis appeared to be reduced in *bKS1Ghr* females (**Supplementary Figure 9H-I**). Hyperphosphorylation of STAT5 in the skeleton and skin was not changed by *Ghr* deletion (**Figure 6I-J**). Failure to normalize weight, skeletal parameters, and skin thickness upon *Ghr* deletion suggests that GHR signaling in skeletal cells and fibroblasts is not required for overgrowth phenotypes. Instead, it supports the model that PDGFRβ^K^ hijacks STAT5 in connective tissue cells, leading to *Igf1* overexpression (**Figure 5B,E**) that potentially mediates overgrowth effects on surrounding tissue. This appears to be the case for all *bKS1^Prrx1Cre^* mice and for male *bKS1^PdgfraCreER^* mice. In female *bKS1^PdgfraCreER^* mice, on the other hand, GHR signaling may generate some basal growth signaling that PDGFRβ^K^ then amplifies to generate overgrowth.

**Figure 6.**
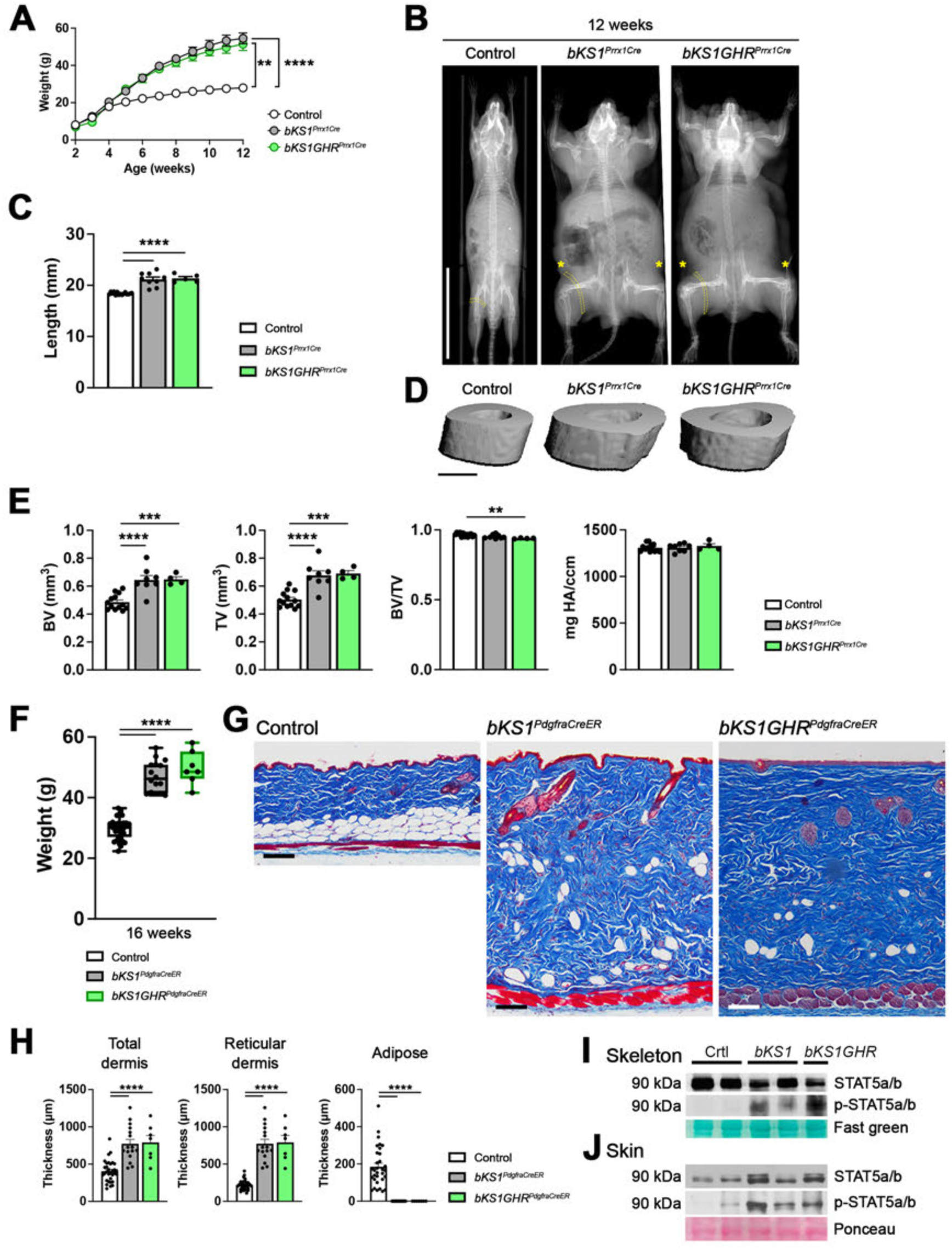
GHR deletion does not rescue overgrowth of male skeleton and skin. **(A)** Weekly body weight measurements of male control, *bKS1^Prrx1Cre^,* and *bKS1GHR^Prrx1Cre^* mice from 2 to 12 weeks old. Control and bKS1 data are the same as Figure 1A. n = 7 for *bKS1GHR^Prrx1Cre^* male mice. **(B)** X-ray images of male control, *bKS1^Prrx1Cre^* and *bKS1 GHR^Prrx1Cre^* mice at 12 weeks old. Dotted yellow bands indicate hindlimb muscle. Yellow asterisks indicate thick skin. Scalebar, 3 cm. (C) Measurements of male tibia length from control, *bKS1^aRCreEM^,* and *bKS1GHR^aRCreEM^* at 12 weeks old. Control and bKS1 data are the same as Figure 2A. (D-E) Micro computed tomography scans and quantifications of male tibia cortical bone from control, *bKS1^Prrx1Cre^,* and *bKS1 GHR^aRCreEM^* mice. Control and bKS1 quantifications are the same as Figure 2C. Scalebar, 0.5 cm. (F) Final body weight measurements of male control, *bKS1^aRCreER^,* and *bKS1 GHR^aRCreER^* mice at 16 weeks old. Control and bKS1 data are the same as Figure 1B. (G-H) Trichrome stain and quantification of male skin from control, *bKS1^aRCreER^,* and *bKS1GHR^aRCreER^* at 16 weeks old. Control and bKS1 quantifications are the same as Figure 4A. Scalebar, 100 pm. (I-J) Western blots of total and phosphorylated STAT5 in control, bKS1, and bKSIGHR skeleton (femurs and tibias) (I) and skin (J). Fast green or ponceau stain shows similar protein loading. Statistical analyses are a mixed-effects model (A) and one-way ANOVA (C, E, F, and H). *, p < 0.05, **, p < 0.01, p < 0.0005, ***, **p <** 0.0001.

### Skeleton and skin overgrowth depend on Igf1-Igf1r signaling

IGF1 acts in almost every tissue and cell type to promote cell proliferation and tissue growth by activating its receptor, IGF1R. To explore the functional contribution of upregulated IGF1 and IGF1R signaling in overgrown *bKS1* mice, we introduced *Igf1^flox^* ^38^ and *Igf1r^flox^* ^39^ to generate *bKS1Igf1^Prrx1Cre^* and *bKS1Igf1r^Prrx1Cre^* for skeletal analysis (**Supplementary Figure 2C**). Across cohorts and sexes, deletion of *Igf1* or *Igf1r* with *Prrx1Cre* led to a normalization of body growth (**Figure 7A, Supplementary Figure 10A**). MicroCT analysis revealed normalization of skeleton, muscle and skin size in *bKS1Igf1^Prrx1Cre^* males at 12 weeks of age, resulting in a relaxed resting posture reminiscent of control and rescued *bKS1S5^Prrx1Cre^* mice (**Figure 7B**). There was partial rescue of these tissues in *bKS1Igf1r^Prrx1Cre^*males (**Figure 7B**). Female *bKS1Igf1^Prrx1Cre^* and *bKS1Igf1r^Prrx1Cre^* mice were completely rescued (**Supplementary Figure 11B)**. In both sexes, *Igf1* deletion normalized tibia length, and cortical bone volume, while *Igf1r* deletion normalized tibia length but reduced cortical bone volume below the level of controls (**Figure 7C-E, Supplementary Figure 11C-E**). Bone mineralization was unchanged across all genotypes. Normalization of bone parameters by *Igf1* deletion suggests that skeletal cell-derived IGF1 is the driver of skeletal overgrowth in *bKS1* mice. We speculate that circulating IGF1, derived from hepatocytes for example, can maintain normal growth rates following skeletal cell deletion of *Igf1*. Overcorrection of bone parameters by *Igf1r* deletion suggests that local and circulating IGF1 converge on skeletal cell IGF1R to regulate growth of the skeleton.

**Figure 7.**
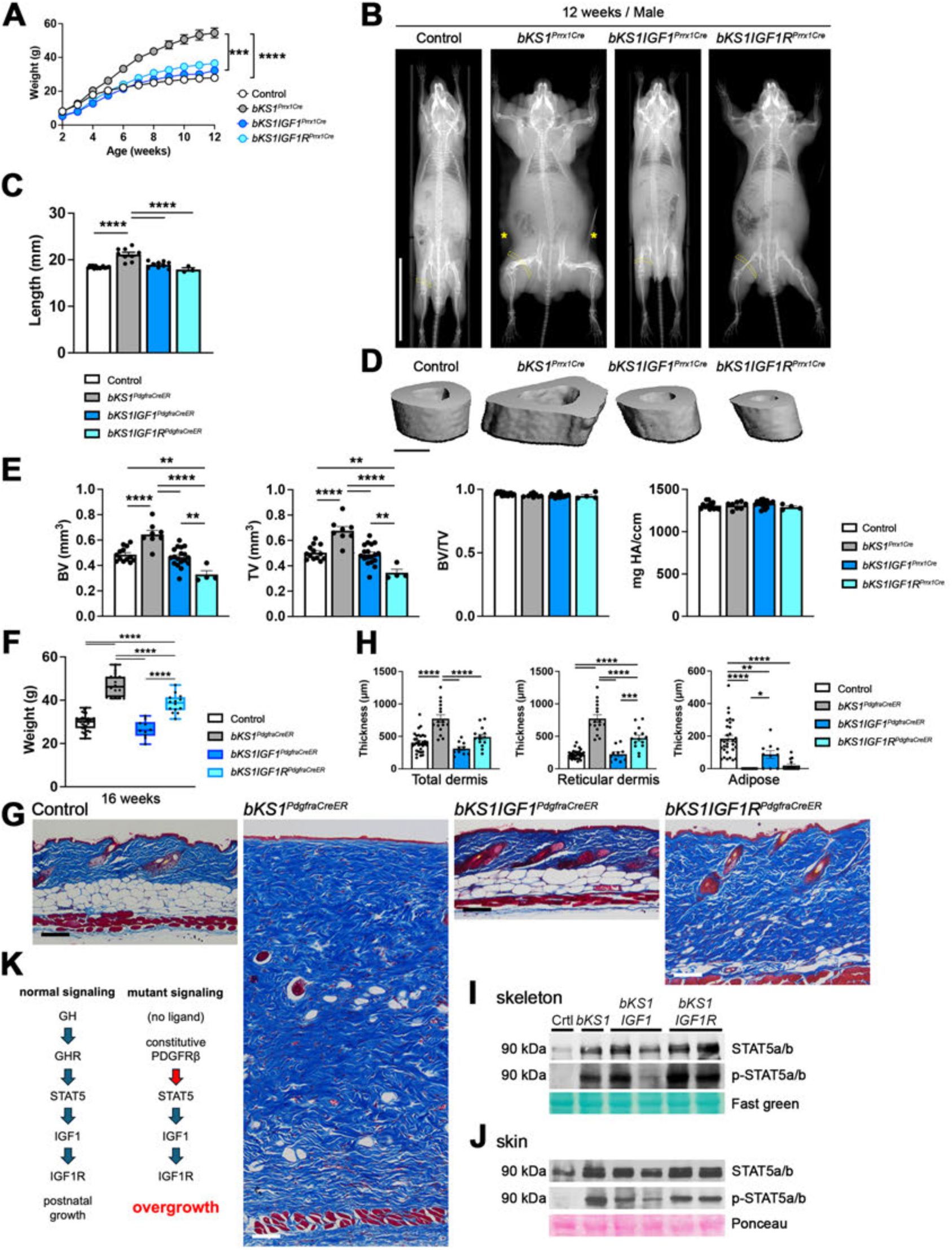
IGF1 and IGF1R deletions rescue overgrowth of male skeleton and skin. **(A)** Weekly body weight measurements of male control, *bKS1^Prrx1Cre^, bKS1IGF1^Prrx1Cre^,* and *bKS1 IGF1 R^Prrx1Cre^* mice from 2 to 12 weeks old. Control and bKS1 data are the same as Figure 1 A. n = 13 for *bKS1IGF1^Prrx1Cre^* and n = 5 for *bKS1 IGF1 R^Prrx1Cre^* male mice. **(B)** X-ray images of male control, *bKS1^Prrx1Cre^, bKS1IGF1^Prrx1Cre^,* and *bKS1IGF1R^Pm1CK^* mice at 12 weeks old. Dotted yellow bands indicate hindlimb muscle, hypertrophied in *bKS1^Prrx1Cre^.* Yellow asterisks indicate thick skin in *bKS1^Prrx1Cre^* Scalebar, 3 cm. **(C)** Measurements of male tibia length from control, *bKS1^Prrx1Cre^*, bKS1IGF1*^Prrx1Cre^*, and *bKS1IGF1R^Prrx1Cre^* at 12 weeks old. Control and bKS1 data are the same as Figure 2A. **(D-E)** Micro computed tomography scans and quantifications of male tibia cortical bone from control, *bKS1^Prrx1Cre^, bKS1IGF1^Prrx1Cre^,* and *bKS1 IGF1 R^Prrx1Cre^* mice. Control and bKS1 quantifications are the same as Figure 2C. Scalebar, 0.5 cm. **(F)** Final body weight measurements of male control, *bKS1^aRCreER^, bKS1IGF1^aRCreER^,* and *bKS1IGF1R^aRCreER^* mice at 16 weeks old. Control and bKS1 data are the same as Figure 1B. (G-H) Trichrome stain and quantification of male skin from control, *bKS1^aRCreER^, bKS1IGF1^aRCreER^,* and *bKS1IGF1R^aRCreER^* at 16 weeks old. Control and bKS1 quantifications are the same as Figure 4A. Scalebar, 100 pm. (I-J) Western blots of total and phosphorylated STAT5 in control, bKS1, bKS1IGF1 and bKS1IGF1R skeleton (femurs and tibias) (I) and skin (J). Fast green or ponceau stain shows similar protein loading. (K) Schematic signaling models for normal postnatal growth and mutant overgrowth. Statistical analyses are a mixed-effects model (A) and one-way ANOVA (C, E, F, and H). *, *p* < 0.05, **, *p <* 0.01, ***, *p <* 0.0005, **’*, *p <* 0.0001.

We also generated *bKS1Igf1^PdgfraCreER^* and *bKS1Igf1r^PdgfraCreER^*for skin analysis (**Supplementary Figure 2E**). Deletion of *Igf1* ligand achieved complete normalization of weight in both sexes. Deletion the receptor achieved complete normalization of female weight, but only partially rescued male weight with *PdgfraCreER* (**Figure 7F, Supplementary Figure 11F**). Histology revealed normalization of skin thickness and adiposity in *bKS1Igf1^PdgfraCreER^* males at 16 weeks of age, resulting in normal thickness of the total skin and upper dermis, and near-complete restoration of dWAT (**Figure 7G-H**). For *bKS1Igf1r^PdgfraCreER^*male skin there was partial rescue, as some mice achieved normalization of skin thickness but without full restoration of dermal fat (**Figure 7G-H**). For females, collagen deposition between adipocytes was rescued in *bKS1Igf1^PdgfraCreER^* and *bKS1Igf1r^PdgfraCreER^* skin compared to *bKS1^PdgfraCreER^* **(Supplementary Figure 11G-H**). Complete normalization of skin thickness by *Igf1* deletion suggests that fibroblast-derived IGF1 is the driver of skin fibrosis and lipodystrophy in *bKS1* mice, with circulating or other sources of IGF1 maintaining normal tissue growth following fibroblast deletion of *Igf1*. As IGF1 levels would still be high in *bKS1Igf1r^PdgfraCreER^* mice, partial normalization of body weight is consistent with elevated IGF1 acting in a paracrine manner to drive overgrowth of tissues that do not experience *PdgfraCreER*-mediated deletion of *Igf1r*, for example muscle and epidermis.

Western blotting for phospho-STAT5 in skeleton or skin (**Figure 7I-J**) showed that it remained highly phosphorylated after deletion of *Igf1* or *Igf1r*. This is consistent with STAT5 acting upstream of IGF1-IGF1R by regulating the expression of *Igf1*. Taken together, these data indicate that IGF1-IGF1R signaling is required for connective tissue overgrowth driven by mutant PDGFRβ.

### Conserved tissue growth signature in human skeleton and skin

To address whether the molecular features identified in mouse skeletal and skin tissues are conserved in humans, we analyzed publicly available human scRNA-seq datasets. Analysis of human bone scRNA-seq data ^40^ identified six major cell populations including osteoblast and bone marrow stromal cell (BMSC) containing skeletal stem and progenitor cells, vascular endothelial cells (VEC), vascular smooth muscle cells (VSMC), hematopoietic and immune cells, and plasma cells (**Figure 8A**). Consistent with our observations in mouse, human osteoblast and BMSC highly expressed *PDGFRB, GHR*, and *IGF1R*, with moderate expression of *STAT5A, STAT5B*, and *IGF1*. In addition, *IGF1R* was broadly expressed across most cell types except plasma cells. For skin, we integrated four publicly available human skin scRNA-seq datasets ^41–44^ (**Figure 8B**). We identified two major fibroblast populations (fibroblast-1 and fibroblast-2), VEC, lymphatic endothelial cells (LEC), mast cells, immune cells, keratinocyte, neural cells, and VSMC. Both types of human fibroblasts highly expressed *PDGFRB* with moderate expression of *GHR, STAT5A, STAT5B* and *IGF1*. *IGF1R* was broadly expressed across multiple cell types. Together, these observations identify connective tissue cells with potential to activate the PDGFRβ-STAT5-IGF1 signaling axis and mediate overgrowth in humans.

**Figure 8.**
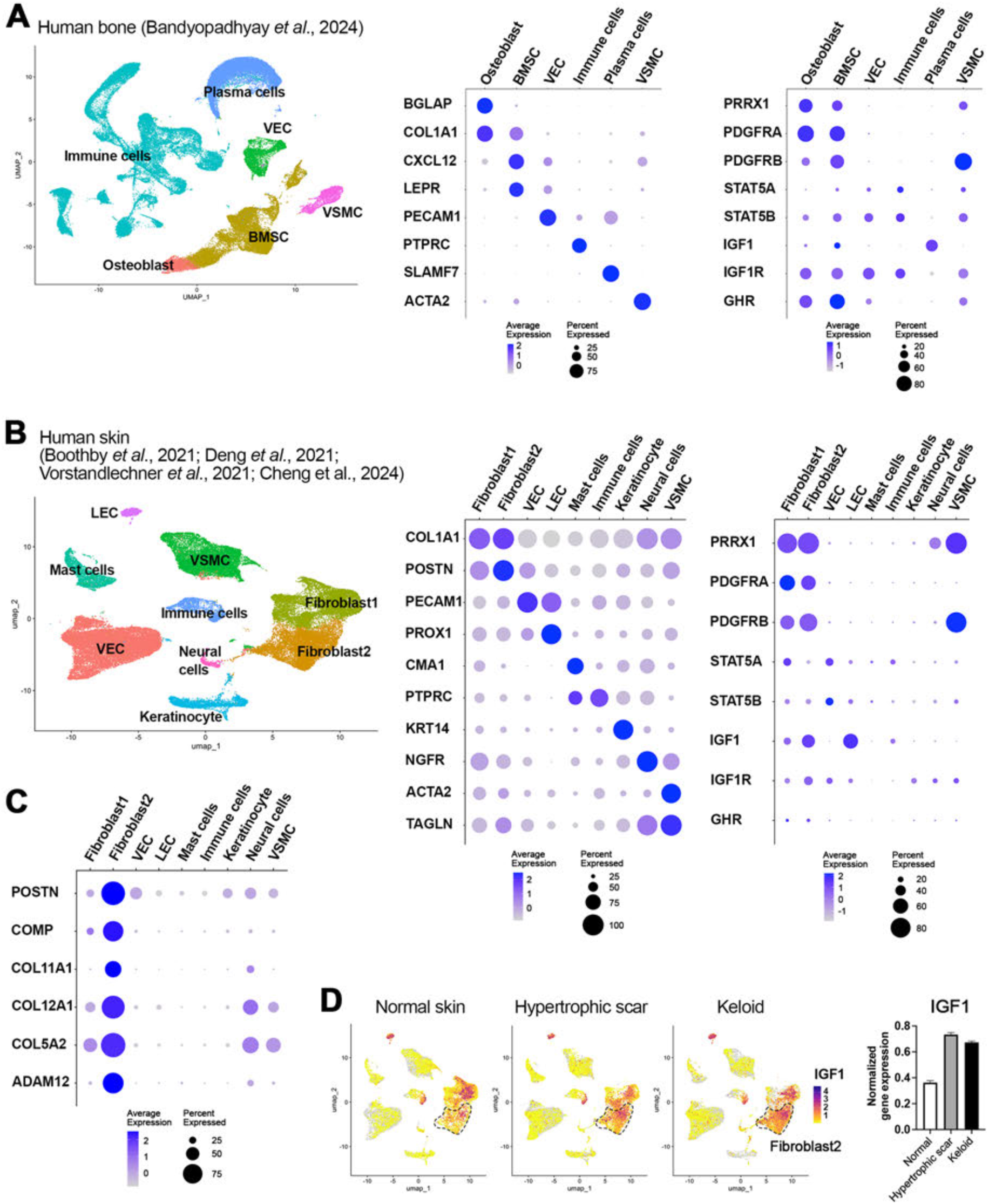
Conserved tissue growth signature in human skeleton and skin. **(A)** UMAP and dotplots of human bone scRNA-seq data from Bandyopadhyay 2024 showing six main cell types (osteoblast. BMSC = bone marrow stromal cells. VEC = vascular endothelial cells. VSMC = vascular smooth muscle cells, immune cells, plasma cells) and marker genes. Rightmost dotplot shows 8 key genes of interest for this study. **(B)** UMAP and dotplots of human skin scRNA-seq data combined from Boothby 2021, Deng 2021, Vorstandlechner 2021, and Cheng 2024 showing nine main cell types (two fibroblast subpopulations. VEC. LEC = lymphatic endothelial cells, mast cells, immune cells, keratinocyte, neural cells. VSMC). Rightmost dotplot shows 8 key genes of interest for this study. (C) Dotplot shows 6 scar-associated fibroblast genes. (D) The expression of IGF1 in normal skin, hypertrophic scar, and keloid. Dotted lines indicate Fibroblast 2 subpopulations. Rightmost bar graph shows normalized expressions of IGF1 in Fibroblast 2 for normal skin (n = 1728 cells), hypertrophic scar (n = 3384 cells), and keloid (n = 6606 cells) with S.E.M.

The integrated skin dataset includes normal skin, hypertrophic scar tissue, and keloid. Scar fibroblasts are characterized by the expression of mesenchymal markers and extracellular matrix proteins ^42,43,45^. In the integrated dataset, the fibroblast-2 cluster was associated with scar pathogenesis, exhibiting high expression of *POSTN, COMP, COL11A1, COL12A1, COL5A2*, and *ADAM12* (**Figure 8C**). Notably, *IGF1* expression was highest in the fibroblast-2 cluster and was enriched in both hypertrophic scar and keloid (**Figure 8B,D**). Considering together the high expression of *Igf1* in mouse fibroblast-2 and human fibroblast-2 clusters (**Supplementary Figure 1C, Figure 8D**) and the dramatic IGF1-dependent skin fibrosis phenotype in *bKS1* mice (**Figure 7G-H**), we suggest an important role for IGF1 in pathological skin fibrosis and keloid in humans.

## DISCUSSION

Normal postnatal growth is governed by pituitary-secreted GH, which circulates as an endocrine hormone. Its binding to GHR on distant cells activates STAT5, leading to expression of IGF1^46–49^. The main finding of this work is that STAT5 and IGF1-IGF1R are essential mediators of overactive PDGFRβ signaling, which operates in osteoblast and fibroblast lineages to cause overgrowth (**Figure 7K**). It is interesting that overgrowth in PDGFRβ-mutant mice at 3-4 weeks old corresponds to the time of GH-dependent growth, which first raised the possibility of crosstalk between PDGF and GH signaling. Our results suggest that the PDGFRβ-STAT5-IGF1 pathway operates largely independently of GH signaling, as deletion of GHR in *Prrx1*- and *Pdgfra-* lineages did not reduce overgrowth in most experiments. Therefore, in cells with mutant PDGFRβ, the STAT5-IGF1 module drives tissue overgrowth. This occurs by direct cell-autonomous mechanisms, exemplified by the expansion of osteoblast and dermal fibroblast lineages that express PDGFRβ^K^, but also by indirect (paracrine) mechanisms leading to expansion of cartilage and muscle where PDGFRβ^K^ is not highly expressed.

The aim of genetic rescue experiments is to test a gene’s functional requirement in the phenotype. Because GHR, STAT5, IGF1, and IGF1R participate in a signaling pathway of central importance for postnatal growth, mutation of these genes has potential to rescue or overcorrect unrelated overgrowth conditions in a non-specific manner. We argue that our study is not a case of non-specific rescue for three reasons. First, we show that *bKS1* mice have elevated STAT5 phosphorylation and *Igf1* expression beyond what is seen in normal growth, indicating that PDGFRβ is mechanistically linked to the pathway. We also used tissue-specific conditional deletion to disrupt the genes in connective tissue cells, which does not affect hepatocytes where STAT5-dependent serum IGF1 originates. Finally, deletion of *Ghr* did not rescue bKS1 overgrowth, but *Stat5ab, Igf1*, and *Igf1r* did rescue, suggesting that PDGFRβ specifically engages the STAT5-IGF1 module without requiring input from GHR.

Mutant PDGFRβ signaling caused skeletal overgrowth with increased length and width of long bones. Bone length is primarily driven by the growth plate, while bone width is mediated by the activity of osteoblasts in periosteum and endosteum. There is strong PDGFRβ expression in the periosteum and endosteum. But expression in the growth plate was restricted to cells of the upper resting zone, and lineage tracing did not suggest a major direct contribution to the growth plate. Cells around the growth plate are known to regulate growth plate activity via secreted signals including parathyroid hormone-related peptide, hedgehog, Wnt, and Fgf ^50–56^. Notably, the growth plate expresses IGF1R and responds to IGF1 stimulation^57–59^. RNA sequencing showed *bKS1^Prrx1Cre^* skeletal upregulation of STAT5 target genes *Igf1* and *Socs2*, which were reversed in *bKS1S5^Prrx1Cre^*bones. Therefore, we suggest that paracrine signaling, especially via IGF1, is likely to mediate PDGFRβ^K^-driven growth plate activity leading to longitudinal growth.

Mutant PDGFRβ signaling caused keloid-like fibrosis with deposition of thicker collagen bundles and obliteration of dermal fat. Lineage tracing revealed that the intersection of *PdgfraCreER* and *bK^Flp^*led to PDGFRβ^K^ expression in fibroblasts of the reticular dermis and dermal adipose tissue. This cell population expanded in fibrotic *bKS1^PdgfraCreER^* skin but was normalized by deletion of *Stat5ab*, *Igf1*, or *Igf1r*. Tissue-level changes were accompanied by increased expression of collagens, cell cycle genes and Wnt signaling. Wnt/β-catenin signaling in the skin is strongly pro-fibrotic^60,61^. Like the skeleton, the skin showed *bKS1^PdgfraCreER^* upregulation of *Igf1* and *Socs2*, which were reversed in *bKS1S5^PdgfraCreER^* skin. Interestingly, *bKS1^PdgfraCreER^*skin also exhibited a striking STAT5-dependent upregulation of hair keratin-associated genes, suggestive of paracrine signaling between the dermal compartment and hair follicles. This might be explained by upregulation of IGF1 or Wnt signaling, as both are known to act on hair follicle keratinocytes to promote hair growth^62–64^. Skin fibrosis exhibited significant sexual dimorphism in *bKS1^PdgfraCreER^* mice. This sex effect is likely related to the postnatal timing or mosaicism of *PdgfraCreER*, as *bKS1* mice generated with a germline Sox2Cre did not exhibit sexually dimorphic skin fibrosis^15^.

GHR activates STAT5 via Janus kinase, but PDGFRβ can activate STAT5 even in the presence of Janus kinase inhibitors^65^. Our experimental approach treated STAT5a and STAT5b as a single entity. Which STAT5 is more important to the overgrowth phenotype? Global *Stat5a/b* double knockout significantly reduces murine growth^66,67^. When disrupted separately, *Stat5b* moderately reduces growth, especially in males, but *Stat5a* deletion alone does not^66,68^. This suggests that both proteins are involved in growth control, with some redundancy, and STAT5b is particularly responsible for sexually dimorphic growth in male mice. We can therefore speculate that both proteins are involved in PDGFRβ-driven overgrowth and STAT5b may be important for male-specific skin fibrosis. The consequence of *Stat5a/b* deletion in the *Prrx1Cre* or *PdgfraCreER* lineages of wild type mice remains uncharacterized, but the tendency to overcorrect bone phenotypes supports current understanding that STAT5 is essential for normal bone growth.

Mutations leading to PDGFRβ activation have been linked to three human diseases: Kosaki overgrowth syndrome (KOGS), Penttinen syndrome (PS), and myofibromatosis. The molecular basis of how different mutations lead to different diseases is unclear, but it is reasonable to speculate that mutation-specific signaling creates distinctive phenotypes. The mice in the current study have a PDGFRβ kinase domain mutation (D849V) corresponding to the human D850V variant associated with sporadic myofibromatosis^6,14^. Drastic connective tissue overgrowth in these mice, on a *Stat1^-/-^*genetic background^15^, initially led us to explore mechanisms of overgrowth. It is important to point out that KOGS *per se* has been associated with juxtamembrane domain mutations P584R and W565R rather than kinase domain mutations. We recently generated a more precise KOGS mouse model with a P583R mutation corresponding to human P584R^65^, and another group has characterized a W565R model^69^. Both models result in cranial bone overgrowth, and the P583R model additionally exhibits skeleton and skin overgrowth on a *Stat1^-/-^* background^65^. Expression of PDGFRβ and components of the STAT5-IGF1-IGF1R axis are conserved in human tissue. We therefore speculate that signaling mechanisms described here will be conserved in other PDGFRβ overgrowth mouse models and in humans.

## MATERIALS AND METHODS

### Animals

All animal experiments were performed according to procedures approved by the Institutional Animal Care and Use Committee at the Oklahoma Medical Research Foundation. Mice were maintained on a 12 hr light/dark cycle and housed in groups of two to five with unlimited access to food and water. All strains were maintained on a mixed C57BL6/129 genetic background at room temperature. Both males and females were analyzed. The following strains were utilized: *Prrx1Cre* (MGI:2450929), *PdgfraCreER^T2^* (MGI:6201731)*, Sox2Cre* (MGI:2656539)*, Stat1^flox^* (MGI:4821871), *Stat5a/b^flox^* (MGI:3055318+MGI:3055319), *Igf1^flox^* (MGI:2152423), *Igf1r^flox^*(MGI:2389580), *Ghr^flox^* (MGI:5550361), *R26-FSF-tdTomato* (MGI:6260212), and *Pdgfrb^Flp^* (MGI:6728540). *Pdgfrb^KFlp^* was generated by the Olson lab ^32^. Breeding strategies are shown in Supplementary Figure 2. For experiments involving PDGFRaCreER^T2^, mice received one gavage of 100 mg/kg tamoxifen after genotyping between P10-P14.

### Western blot

For skeletal tissues, femurs and tibias were harvested at 4-8 weeks of age and bone marrow was flushed out. Bones were frozen in liquid nitrogen and were pulverized with a mortar and pestle. For skin tissues, dorsal skins at 16 weeks of age were harvested, hair was removed using depilatory cream, frozen in liquid nitrogen, and pulverized with a tissue pulverizer. The homogenized tissues were dissolved in RIPA buffer (50mM Tris pH 7.4, 1% NP-40, 0.25% sodium deoxycholate, 150 mM NaCl, 0.1% sodium dodecyl sulfate) with 1 mM NaF, Na3VO4, PMSF, and 1x protease inhibitor cocktail (Complete, Roche). After determining protein concentration with Pierce BCA assay, proteins were heated with 4X sample buffer with 5% beta-mercaptoethanol at 70 °C for 5 minutes for skeletal samples or at 95 °C for 5 minutes for skin samples. 40-100 μg of proteins separated by SDS-PAGE and transferred to nitrocellulose membrane. The membranes were blocked with 5% BSA, incubated with primary antibodies overnight at 4 °C, probed with horseradish peroxidase (HRP)-conjugated secondary antibody (1:5000, Jackson ImmunoResearch). Blots were developed with Pierce ECL Western blotting substrate (ThermoFisher) and autoradiography film (Santa Cruz). Fast green or ponceau S were used to stain proteins on membranes post-transfer.

### Histology and Immunostaining

For histological analysis, tibias were harvested at 3 weeks of age and fixed with 10% neutral-buffered formalin (NBF) overnight at RT. Fixed skeletal tissues were decalcified in 0.5 M EDTA (pH 7.4) for 1-2 days, embedded in paraffin, sectioned at 6 μm, and stained with hematoxylin and eosin (H&E) or safranin O. For skin, hair was removed using depilatory cream at 16 weeks of age, followed by fixation as above. Fixed skin tissues were embedded in paraffin, sectioned at 8 μm, and stained with Trichome stain or picrosirius red. The stains were imaged using a Nikon Eclipse 80i microscope with a digital camera. Birefringence of picrosirius red stains were imaged with polarized light.

For lineage tracing and immunostaining, whole legs containing femurs, knee joint, and femurs were harvested at 3 weeks of age and fixed with 4% paraformaldehyde overnight at 4 °C. Fixed skeletal tissues were decalcified in 0.5 M EDTA (pH 7.4) for 1-2 days overnight at 4 °C, dehydrated with 20% sucrose in PBS, and cryosectioned at 100 μm thickness. Skin was harvested at 16 weeks and fixed with 4% paraformaldehyde overnight at 4 °C. Fixed skin was dehydrated in 20% sucrose in PBS and cryosectioned at 20 μm thickness. The sectioned tissues were stained with primary antibodies, conjugated secondary antibodies (Jackson ImmunoResearch), and DAPI (Sigma) for nuclei stain. Primary antibodies used in this manuscript include antibodies for Endomucin (Santa Cruz, sc-65495, 1:200), PDGFRbeta (R&D, AF1042, 1:200), Perilipin-1 (Cell Signaling, 9349, 1:250), and Vimentin (Cell signaling, 5741, 1:200). Imaging was performed with a Nikon C2 confocal for skeleton and a Nikon Eclipse 80i microscope for skin.

### X-ray microCT

Mouse whole body X-ray images were captured shortly after the euthanasia using a vivaCT 40 microCT (Scanco Medical). Tibias at 12 weeks of age were fixed with 10% NBF for 3-4 days and scanned with the microCT at 70kVp and 110uA. Tibial cortical bones right above tibio-fibular junction were analyzed to measure bone volume (BV, mm^3^), total volume (TV, mm^3^), BV/TV, and mean density (mg HA/ccm).

### RNA sequencing and analysis

Femurs and tibias in controls, *bKS1^Prrx1Cre^*, and *bKS1S5^Prrx1Cre^* at 3-6 weeks of ages were harvested and bone marrow were flushed out. The bones were frozen in liquid nitrogen in the mortar and ground with pestle. Dorsal skins in controls, *bKS1^PdgfraCreER^*, and *bKS1S5^PdgfraCreER^* at 16 weeks of age were harvested, frozen in liquid nitrogen, and pulverized with a tissue pulverizer. The homogenized skeletal and skin tissues were dissolved in Trizol to isolate RNA, followed by sequencing. Raw sequencing reads were aligned to the mm10 mouse reference genome and quantified using Spliced Transcripts Alignments to a Reference (STAR, v2.7.10b) ^70^ and featureCounts (v 2.0.8). Read counts were normalized and processed using R (v4.4.1) and DESeq2 ^71^. Differentially expressed genes (DEGs) were determined by p-value ≤ 0.05 and an absolute log2 fold change ≥ 0.25. DEGs were further analyzed with the Database for Annotation, Visualization and Integrated Discovery (DAVID) and gene set enrichment analysis (GSEA, M2 curated gene sets) to identify gene ontology categories and KEGG pathways ^72,73^. Heatmaps were plotted using pheatmap, an R package ^74^. To quantify gene-set expression, variance-stabilized expression values were standardized for each gene across samples by Z-score transformation. Mean Z-scores were calculated across all genes within each indicated upregulated or downregulated gene category.

### Single-cell RNA sequencing data analysis

Single-cell RNA sequencing (scRNAseq) data were collected from GSE128423 (mouse skeleton) ^21^, GSE156636 (mouse skeleton) ^22^, GSE153596 (mouse skin) ^23^, GSE253355 (human skeleton) ^40^, GSE183031 (human normal skin) ^41^, GSE156326 (human normal skin and hypertrophic scar) ^43^, GSE163973 (human hypertrophic scar and keloid) ^42^, GSE243716 (human hypertrophic scar and keloid) ^44^. The datasets were analyzed using R (v4.4.1) and Seurat 4.4.0 ^75^. Clusters were visualized using uniform manifold approximation and projection (UMAP) and defined with cell type markers. Dot plots were used to show the expression levels of cell type markers and 8 target genes in this study (*Prrx1*, *Pdgfra*, *Pdgfrb*, *Stat5a*, *Stat5b*, *Igf1*, *Igf1r*, and *Ghr*). scCustomize was used to visualize IGF1 expression ^76^.

### Quantification and statistical analysis

ImageJ (National Institute of Health) ^77^ and NIS-Elments D 3.2 (Nikon) were used to quantify safranin O-stained cartilage (area and length), H&E-stained growth plate (resting zone, proliferating zone, and hypertrophic zone), tdTomato-labeled growth plate (tomato area ratio), tdTomato-labled BMSCs (tomato area ratio), Trichome-stained skin thickness (thickness), and tdTomato-labeled skin area (tomato area ratio). Area and area ratio were measured using the polygon selection tool, and length and thickness were measured using the straight tool. Body weights were measured at every week from 2 to 12 weeks for Prrx1Cre strains and at the 16 weeks endpoint for PDGFRaCreER^T2^ strains in both sexes. Tibia lengths were measured at 12 weeks. Statistical calculations were performed using GraphPad Prism 10 with unpaired two-tailed *t-test*, one-way ANOVA (turkey’s multiple comparisons), or a mixed-effects model (Sidak’s multiple comparisons) representing with mean ± standard error of the mean (SEM).

## Data availability

RNA sequencing data generated in the current study from skeletal and skin tissues will be deposited in the Gene Expression Omnibus public repository. The submission is currently in progress.

## Author contributions

H.R.K and L.E.O conceived and designed the study. H.R.K and L.E.O performed all the experiments and analysis. A.R contributed to mouse colony management and histology. H.R.K wrote the manuscript. H.R.K and L.E.O edited the manuscript. L.E.O supervised the research.

## Acknowledgements

H.R.K. was supported by NIH/NHLBI Ruth L. Kirschstein NRSA F32-HL142222. This research was supported by R01-AR073828 and R01-AR080896 (L.E.O), by the Oklahoma Center for Adult Stem Cell Research (a program of TSET), and by grants from the Oklahoma City-based Presbyterian Health Foundation to L.E.O.

---

**Supplementary Figure 1.**
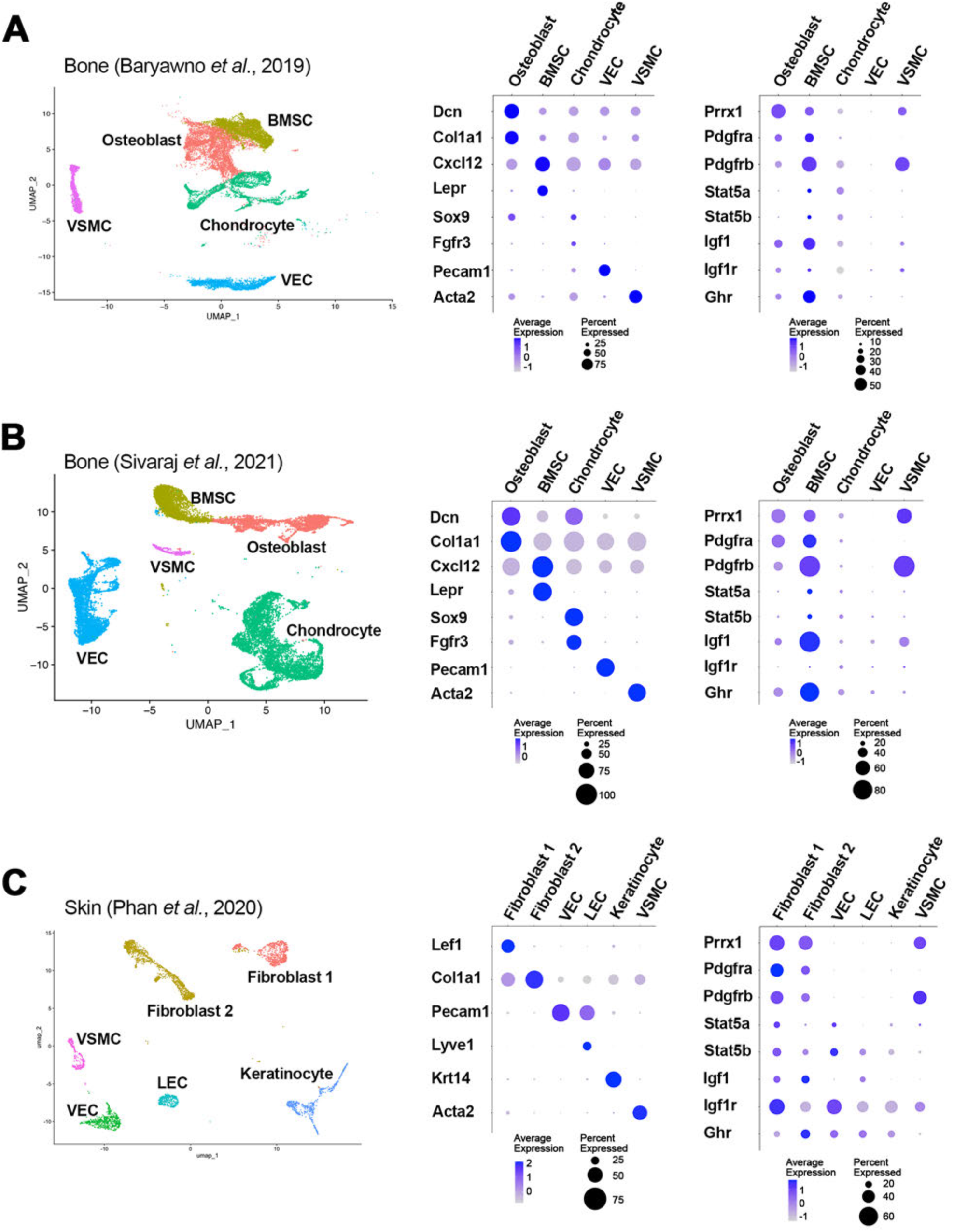
Re-analysis of single-cell RNA sequencing data from skeleton and skin. **(A-B)** UMAP and dotplots of bone scRNA-seq data from Baryawno 2019 and Sivaraj 2021 showing five main cell types (BMSC = bone marrow stromal cell, VEC = vascular endothelial cell, VSMC = vascular smooth muscle cell) and marker genes. Rightmost dotplot shows 8 key genes of interest for this study. **(C)** UMAP and dotplots of skin scRNA-seq data from Phan 2020 showing four main cell types and marker genes. Rightmost dotplot shows 8 key genes of interest for this study.

**Supplementary Figure 2.**
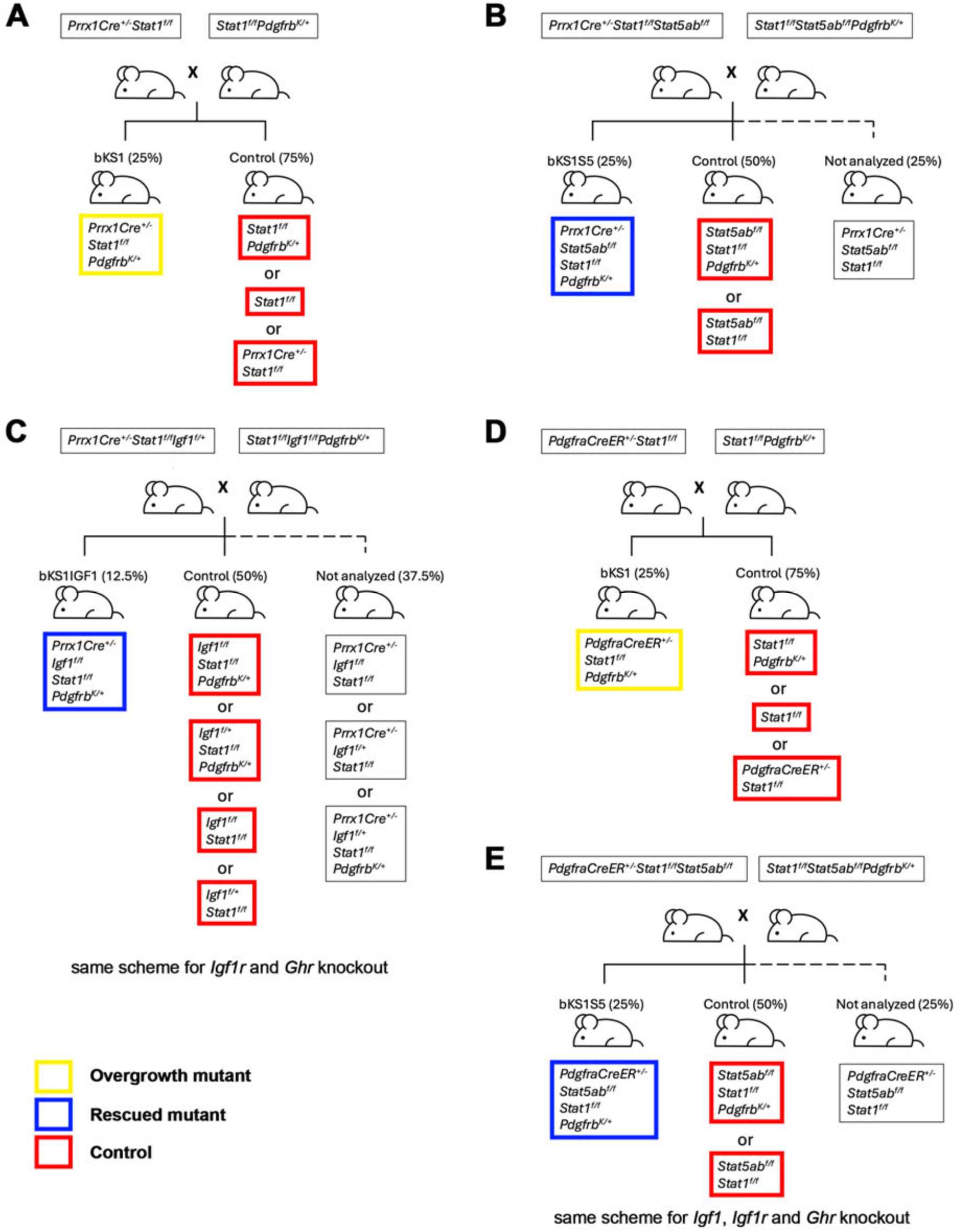
Breeding schemes to activate PDGFRβ^K^ and delete *Statl, Stat5ab, Igfl, Igflr,* and *Ghr* in skeleton and skin. **(A)** Breeding scheme to activate PDGFRp^K^ and delete *Statl* with *Prrx1-Cre* in *bKS1^Prrx1Cra^* mice. **(B)** Breeding scheme to activate PDGFRp^K^ and delete *Statl* and *Stat5ab* with *Prrx1-Cre* in *bKS1S5^Pm1Cra^* mice. **(C)** Breeding scheme to activate PDGFRp^K^ and delete *Statl* and *Igf1* with *Prrx1-Cre* in *bKIGF1^Pm,1Cra^* mice. A similar scheme was used to generate *bKS1IGF1R^Prrx1Cre^* and *bKS1GHR^Prrx1Cre^* mice. **(D)** Breeding scheme to activate PDGFRp^K^ and delete *Statl* with *PdgfraCreER* in *bKS1^aRCraER^* mice. **(E)** Breeding scheme to activate PDGFRp^K^ and delete *Statl* and *Stat5ab* with *PdgfraCreER* in *bKS1S5^aRCraER^* mice. A similar scheme was used to generate *bKS1IGF1^aRCraER^, bKS1IGF1R^aRCraER^,* and *bKS1GHR^aRCraER^* mice.

**Supplementary Figure 3.**
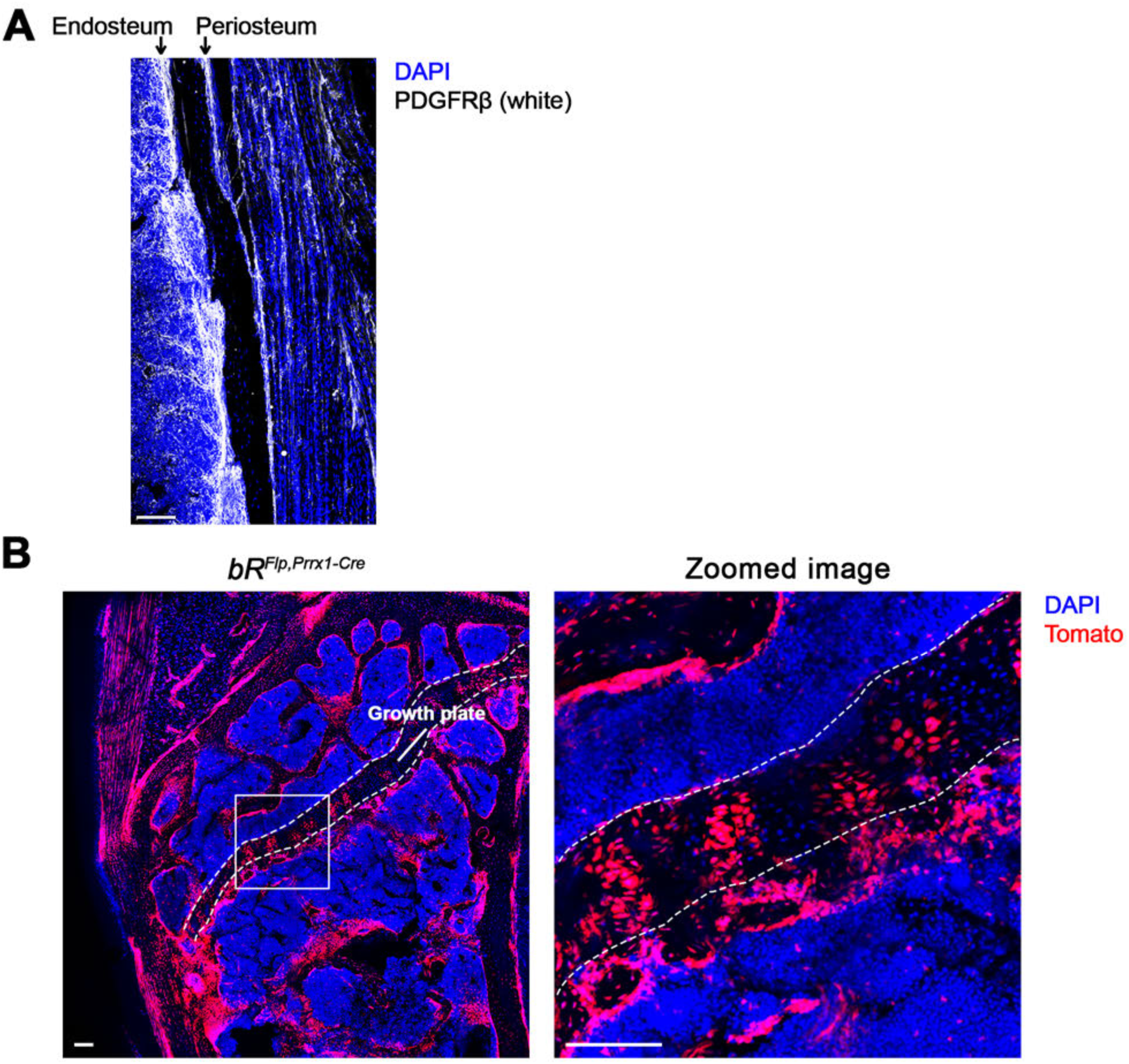
Staining and intersectional lineage tracing of PDGFRp in *Prrx1-Cre.* **(A)** Immunostaining of PDGFRp in diaphysis of femur at 3 weeks old in control mice, with expression in both endosteum and periosteum. Scalebar, 100 pm. **(B)** Confocal microscopy of proximal tibia of *bR^Flp, Prxx1Cre^* at 6 months old. Tomato labeling indicates cells expressing bR^Flp^ including columns of growth plate chondrocytes. Scalebar, 100 pm.

**Supplementary Figure 4.**
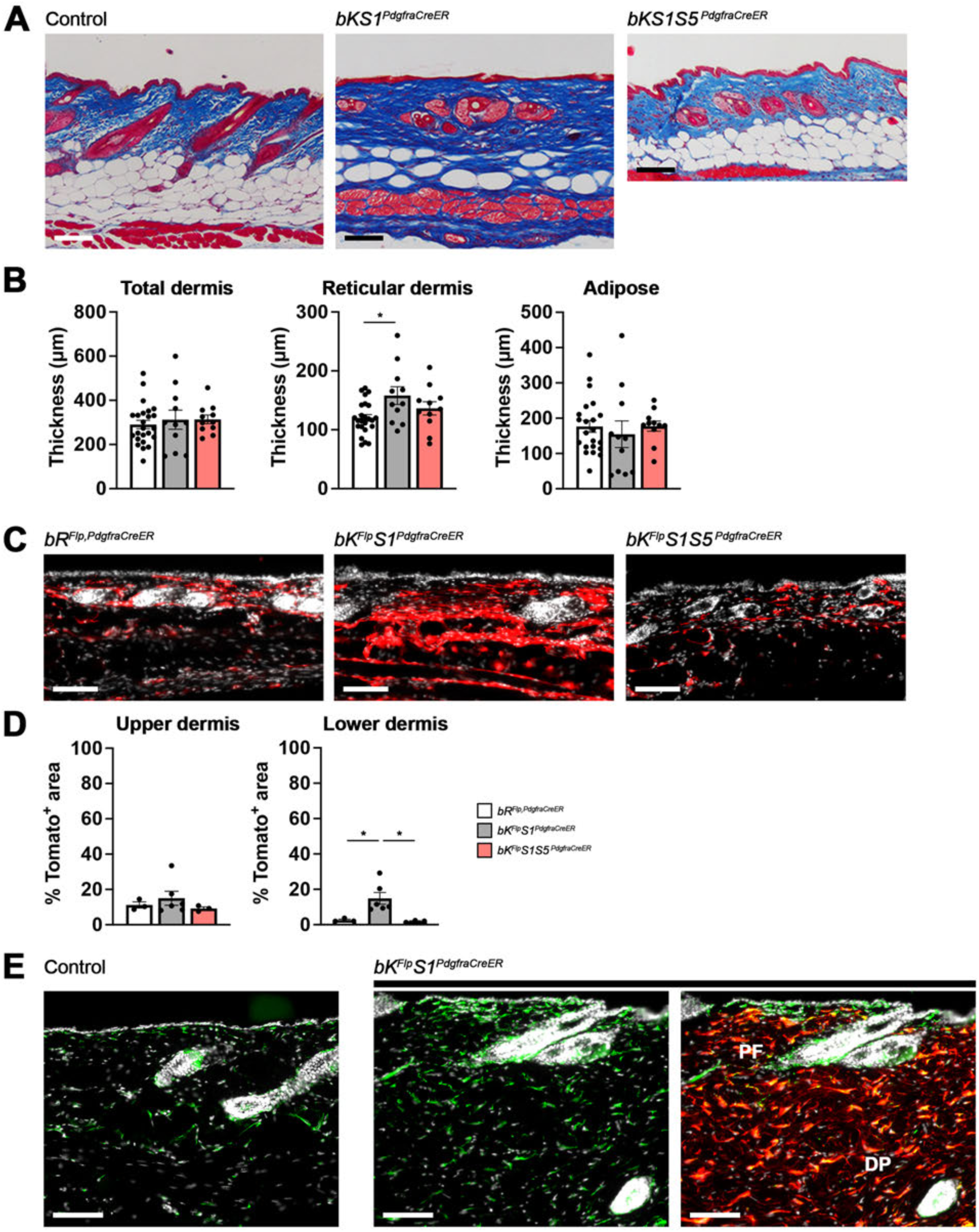
Female dermal fibrosis depends on STAT5. **(A-B)** Trichrome stain of female skin from control, *bKS1^PdglraCreER^,* and *bKS1 S5^Pd9lraCraER^* mice at 16 weeks old. Quantifications of thickness of total dermis, reticular dermis, and dermal white adipose tissue are shown. Scalebar, 100 pm. **(C-D)** Fluorescent images of female skins. Tomato labeling indicates intersectional lineage tracing of *PdgfraCreER* and *bF^Flp^* or *bK^Flp^.* Scalebar, 100 pm. Quantification of Tomato* area in upper and lower dermis are shown. **(E)** Immunofluorescent images of male skin from control and *bKS1^Prrx1Cre^* stained for fibroblast marker Vimentin (green). Tomato labeling indicates intersectional lineage tracing of *PdgfraCreER* and *bK^Flp^* in Vimentin* fibroblasts. Scalebar, 100 pm. Statistical analyses are ANOVA. *, p < 0.05.

**Supplementary Figure 5.**
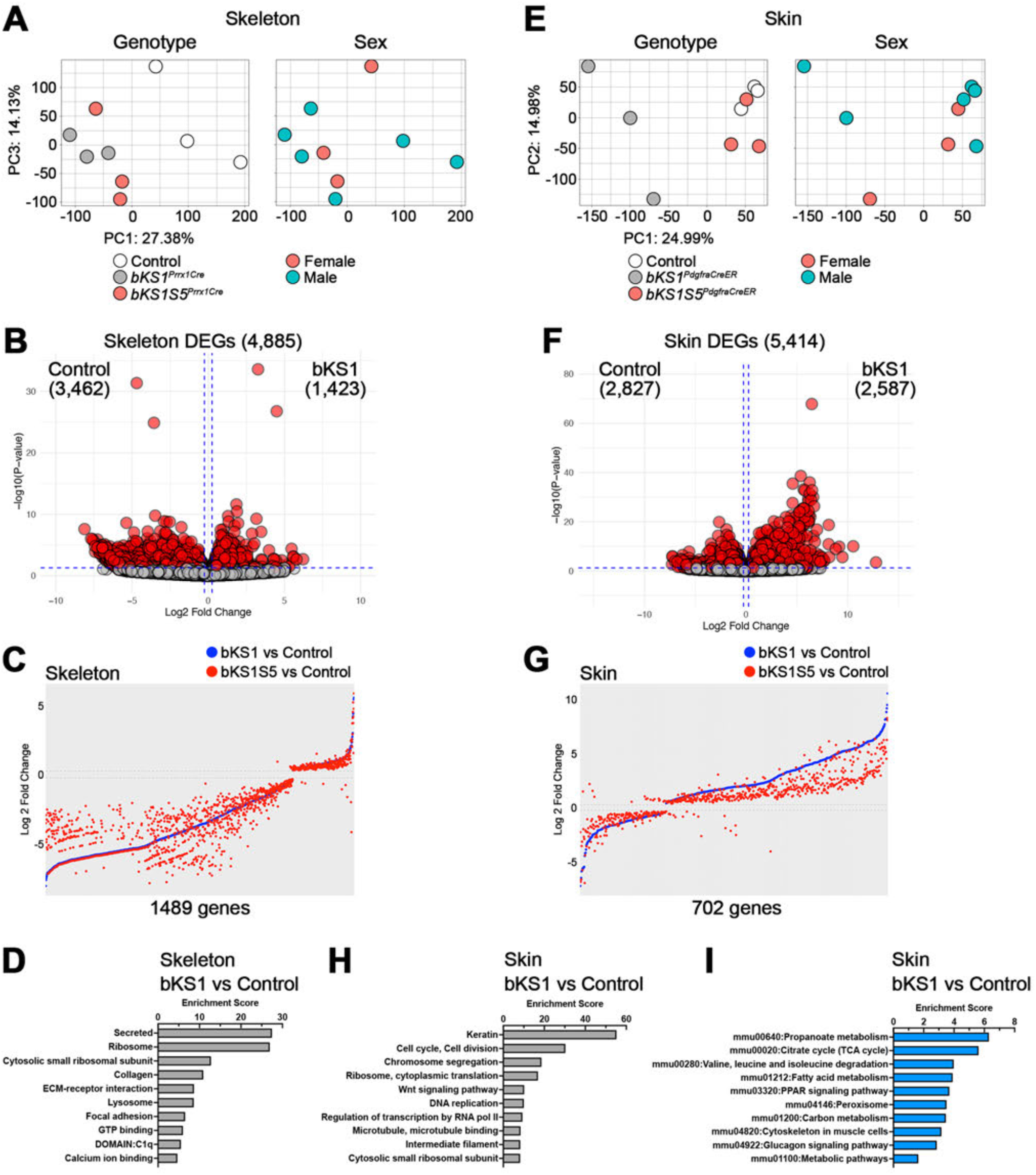
STAT5 deletion rescues gene expression changes in skeleton and skin. **(A)** PCA plots of femur and tibia gene expression from control, *bKS1^Prrx1Cre^,* and *bKS1S5^Prrx1Cre^* mice at 6 weeks old. **(B)** Volcano plot visualizing skeleton differentially expressed genes (DEGs) between controls and *bKS1^Prrx1Cre^* skeletons. **(C)** Comparison of Iog2 fold change of 1489 DEGs shared by *bKS1^Prrx1Cre^* and *bKS1S5^Prrx1Cre^* with 49.6% of the DEGs showing decreased fold change in *bKS1S5^Prrx1Cre^.* **(D)** Top 10 gene ontology categories upregulated in *bKS1^Pmi1Cra^* skeleton compared to control. **(E)** PCA plots of skin gene expression at 16 weeks old. **(F)** Volcano plot visualizing skin DEGs between controls and *bKS1^Pm1Cra^.* **(G)** Comparison of Iog2 fold change of 702 DEGs shared by *bKS1^aRCraER^* and *bKS1 S5^aRCreER^* with 88.2% of the DEGs showing decreased fold change in *bKS1 S5^aRCreER^* skins. (H) Top 10 gene ontology categories upregulated in *t>KS1^Prrx1CreER^* skin compared to control. (I) Top 10 KEGG categories downregulated in *bKS1^aRCreER^* compared to control.

**Supplementary Figure 6.**
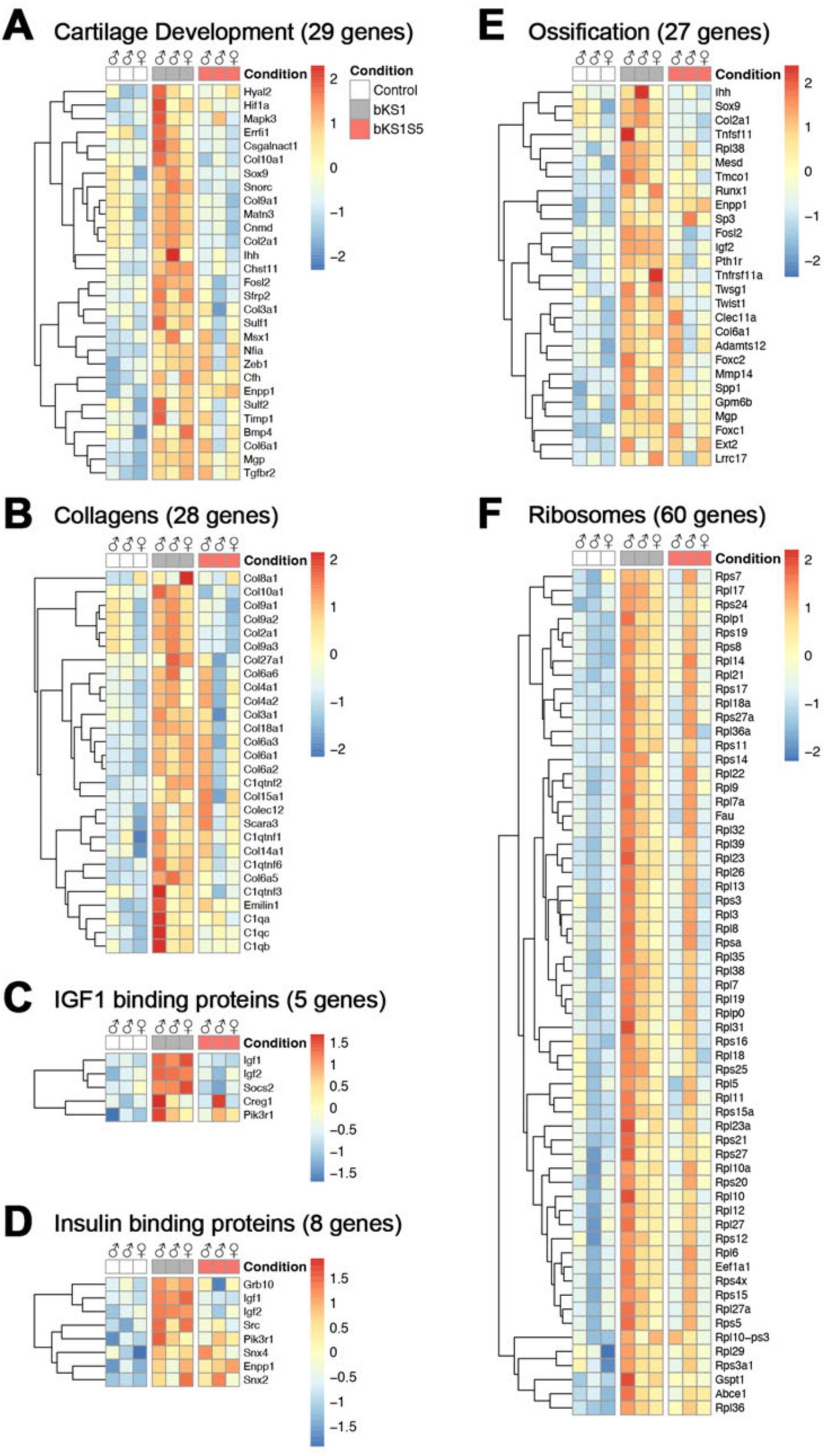
STAT5 deletion rescues gene expression changes in skeleton. Heatmaps indicating pathways and their corresponding genes upregulated in *bKS1^Prrx1Cre^* and normalized in *bKS1S5^Prnl1Cra^*.

**Supplementary Figure 7.**
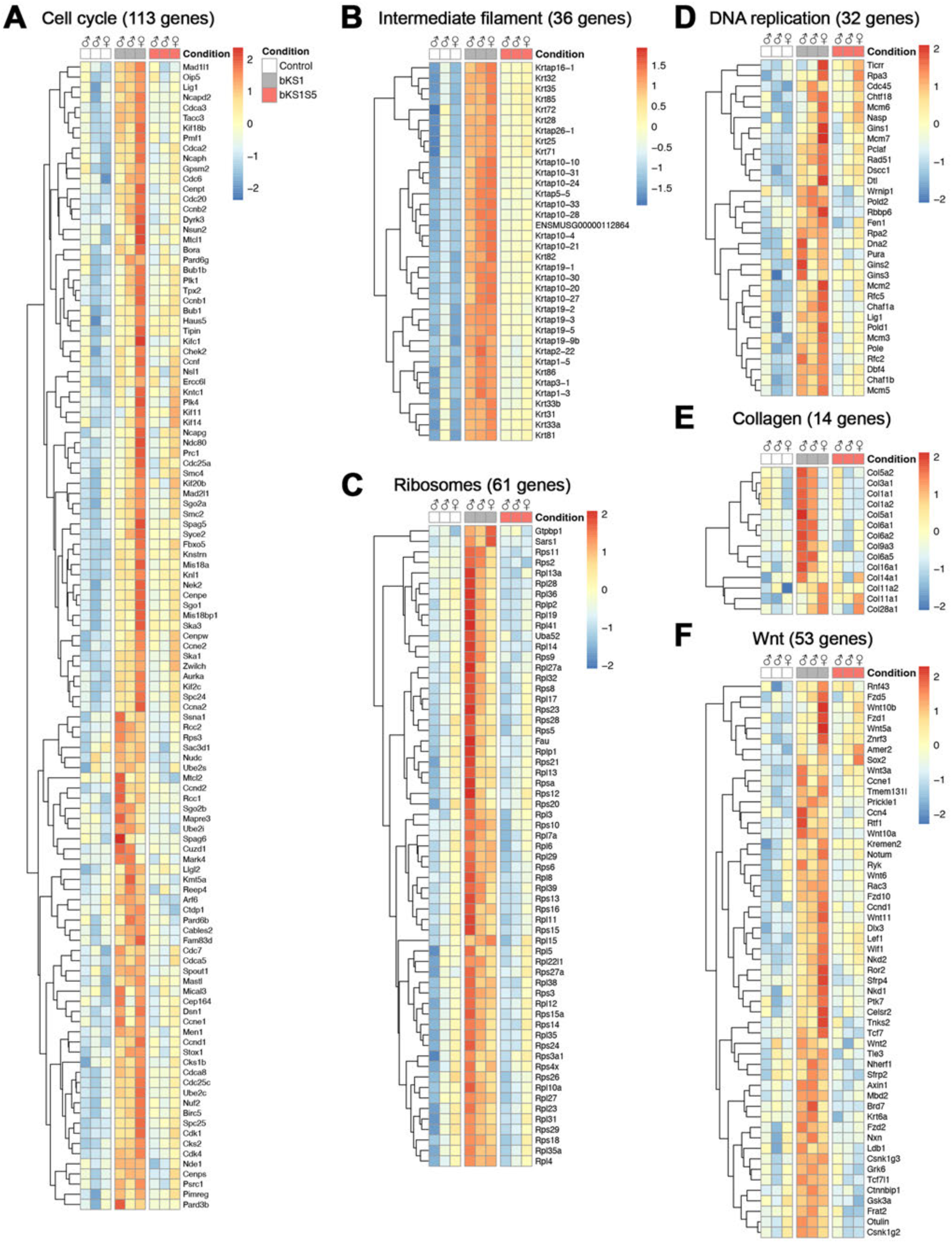
STAT5 deletion rescues gene expression upregulated in bKS1 skin. Heatmaps indicating pathways and their corresponding genes upregulated in *bKS1^aRCreER^* and normalized in *bKS1S5^aRCreER^*.

**Supplementary Figure 8.**
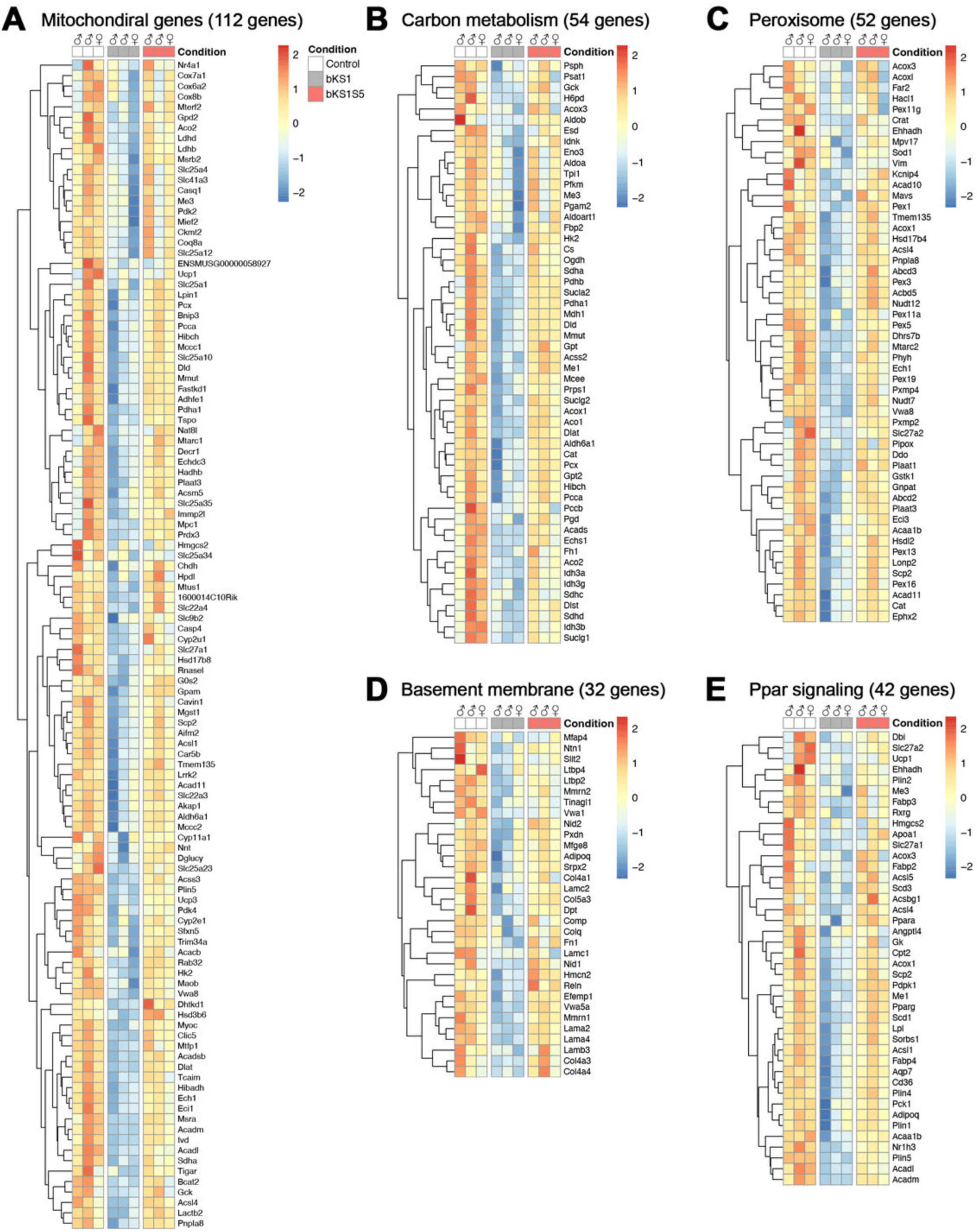
STAT5 deletion rescues gene expression downregulated in bKS1 skin. Heatmaps indicating pathways and their corresponding genes downregulated in *bKS1^aRCreER^* and normalized in *bKS1S5^aRCreER^*.

**Supplementary Figure 9.**
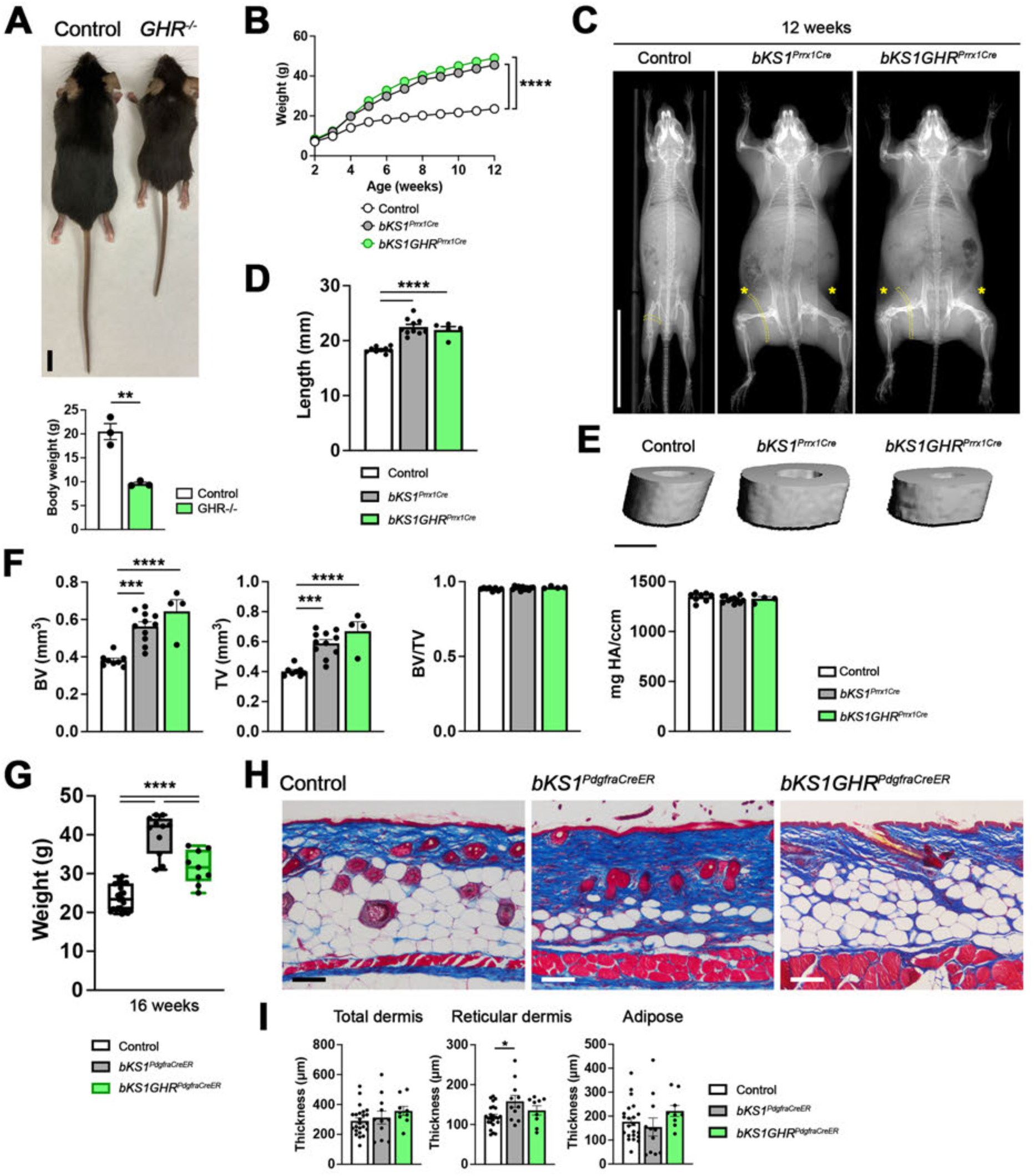
GHR deletion does not rescue overgrowth of female skeleton but partially rescues female skin. **(A)** Photography and body weight measurements of male control and Ghr^flox/flox^Sox2Cre mice at 7 weeks old. n = 6 for *bKS1GHR^Prrx1Cre^* female mice. **(B)** Weekly body weight measurements of female control, *bKS1^Prrx1Cre^,* and *bKS1GHR^Prrx1Cre^* mice from 2 to 12 weeks old. Control and bKS1 data are the same as Figure 1A. **(C)** X-ray images showed 12-week-old female body growth of control, *bKS1^Prrx1Cre^,* and *bKS1GHR^Prrx1Cre^.* Yellow asterisks pointed abdominal skin overgrowth out. Dotted yellow bands highlighted thigh thickness. Scalebar, 3 cm. **(D)** Measurements of female tibia length from control, *bKS1^Prrx1Cre^,* and *bKS1GHR^Prrx1Cre^* at 12 weeks old. Control and bKS1 data are the same as Figure 2A. **(E-F)** Micro computed tomography scans and quantifications of female tibia cortical bone from control, *bKS1^Prrx1Cre^,* and *bKS1GHR^Prrx1Cre^* mice. Control and bKS1 quantifications are the same as Figure 2C. Scalebar, 0.5 cm. **(G)** Final body weight measurements of female control, *bKS1^aRCreER^,* and *bKS1 GHR^aRCraER^* mice at 16 weeks old. Control and bKS1 data are the same as Figure 1B. **(H-l)** Trichrome stain and quantification of female skin from control, *PKS1^Pd9fraCraER^,* and *bKS1GHR^aRCraER^* at 16 weeks old. Control and bKS1 quantifications are the same as Figure 4A. Scalebar, 100 pm. Scalebar, 1 cm. Statistical analyses are unpaired two-tailed *t-test* (A), a mixed-effects model (B), and one-way ANOVA (D, F, G, and I). **, *p < 0.01, p <* 0.0005, “**, *p <* 0.0001.

**Supplementary Figure 10.**
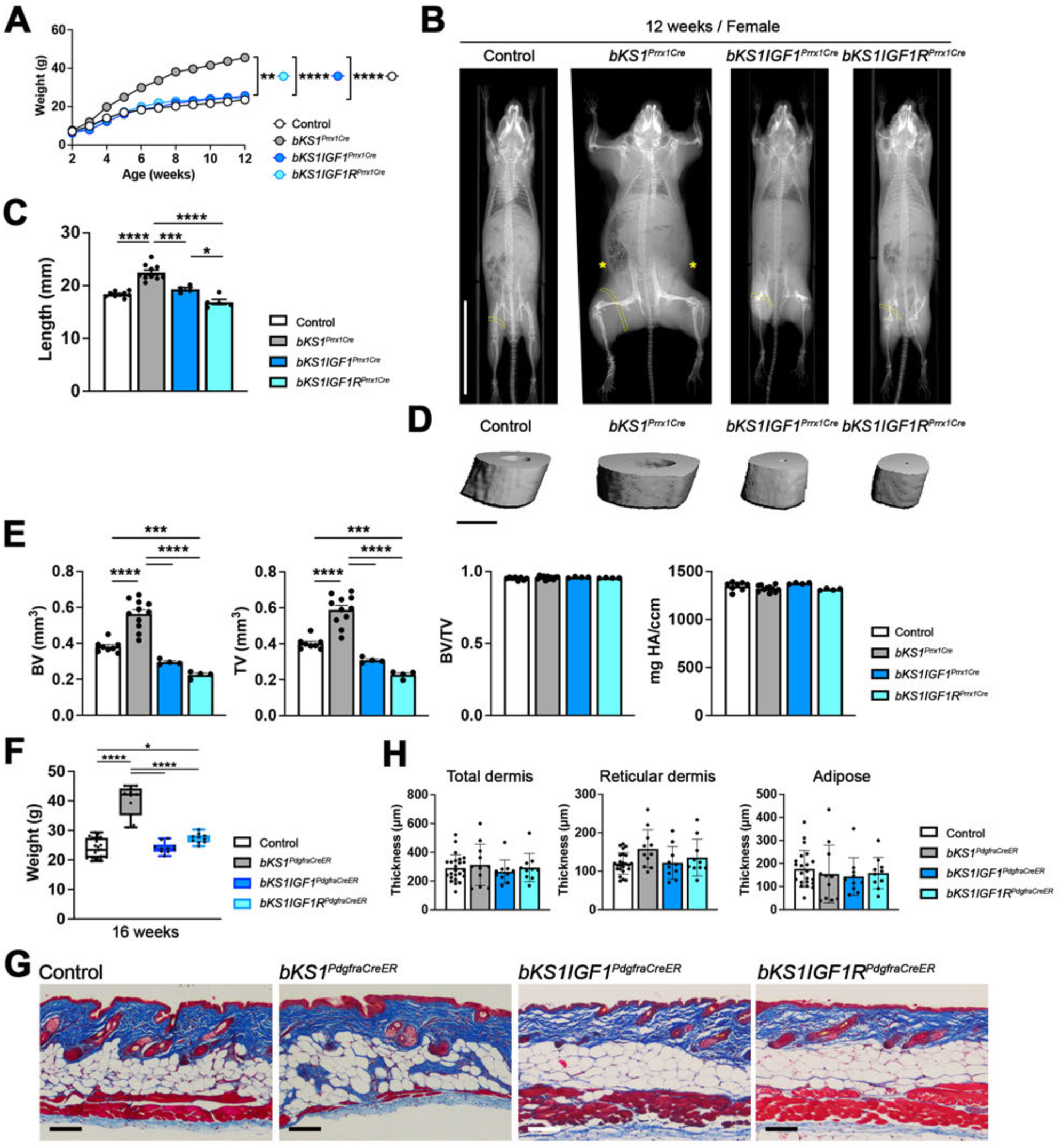
IGF1 and IGF1R deletions rescue overgrowth of female skeleton and skin. **(A)** Weekly body weight measurements of female control, *bKS1^Prrx1Cra^, bKS1IGF1^Prrx1Cre^,* and *bKS1IGF1R^Prrx1Cre^* mice from 2 to 12 weeks old. Control and bKS1 data are the same as Figure 1A. n = 9 for *bKS1IGF1^Prrx1Cre^* and n = 6 for *bKS1IGF1R^Prrx1Cre^* female mice. **(B)** X-ray images of female control, *bKS1^Prrx1Cre^, bKS1IGF1^Prrx1Cre^,* and *bKS1IGF1R^Prrx1Cre^* mice at 12 weeks old. Dotted yellow bands indicate hindlimb muscle, hypertrophied in *bKS1^Prrx1Cre^.* Yellow asterisks indicate thick skin in *bKS1^Prrx1Cre^.* Scalebar, 3 cm. **(C)** Measurements of female tibia length from control, *bKS1^Prrx1Cre^, bKS1IGF1^Prrx1Cre^,* and *bKS1IGF1R^Prrx1Cre^* at 12 weeks old. Control and bKS1 data are the same as Figure 2A. **(D-E)** Micro computed tomography scans and quantifications of female tibia cortical bone from control, *bKS1^Prrx1Cre^, bKS1IGF1^Prrx1Cre^,* and *bKS1IGF1R^Prrx1Cre^* mice. Control and bKS1 quantifications are the same as Figure 2C. Scalebar, 0.5 cm. **(F)** Final body weight measurements of female control, *bKS1^aRCraER^, bKS1IGF1^aRCraER^,* and *bKS1IGF1R^aRCreER^* mice at 16 weeks old. Control and bKS1 data are the same as Figure 1B. **(G-H)** Trichrome stain and quantification of female skin from control, *bKS1^aRCraER^, bKS1IGF1^aRCraER^*, and *bKS1IGF1R^aRCraER^* at 16 weeks old. Control and bKS1 quantifications are the same as Figure 4A. Scalebar, 100 pm. Statistical analyses are a mixed-effects model (A) and one-way ANOVA (C, E, F, and H). *, *p* < 0.05, ***, *p <* 0.0005, *“*, *p <* 0.0001.

## Notes

**Conflict of interest:** The authors have no conflicts of interests.

### Competing Interest Statement

The authors have declared no competing interest.

